# Translocation of gut *Enterococcus faecalis* trains myeloid bone marrow progenitors via the C-type lectin receptor Mincle

**DOI:** 10.1101/2024.11.07.622236

**Authors:** Iñaki Robles-Vera, Aitor Jarit-Cabanillas, Paola Brandi, María Martínez-López, Sarai Martínez-Cano, Cristina González-Correa, Javier Moleón, Juan Duarte, Laura Conejero, Pablo Mata-Martínez, Carmen María Díez-Rivero, Marta Bergón-Gutiérrez, Iván Fernández-López, Manuel J. Gómez, Ana Quintas, Ana Dopazo, Fátima Sánchez-Cabo, Esther Pariente, Carlos del Fresno, José Luis Subiza, Salvador Iborra, David Sancho

## Abstract

Impairment of the intestinal barrier allows the systemic translocation of commensal bacteria, inducing a pro-inflammatory state in the host. To address potential mechanisms underlying this link, we explored innate immune responses following increased gut permeability upon administration of dextran sulfate sodium (DSS) to mice. Microbiota translocation induced trained immunity (TI) in mouse myeloid bone marrow progenitors (BMPs). *Enterococcus faecalis* was the predominant bacteria detected in the bone marrow (BM) following DSS treatment. Notably, live or heat-killed *E. faecalis* induced reprogramming of mouse BMPs *in vitro* and *in vivo* and enhanced macrophage inflammatory activity, also training human monocytes. *E. faecalis* sensing by the C-type lectin receptor Mincle (*Clec4e^−/-^*) is essential for TI induction in BMPs. Consequently, *Clec4e*^−/-^ mice showed impaired TI upon *E. faecalis* or reduced pathology following DSS treatment. Our results identify *E. faecalis* as inducer of TI-related inflammation that may contribute to pathologies associated with increased gut permeability.

## Introduction

The host immune system is exposed and has coevolved with trillions of microorganisms, drawing a fine balance between the development of a competent immune system that promotes inflammatory responses to eliminate pathogens while maintaining the symbiotic relationship with commensal microbiota^1^. However, the imbalance in gut microbial composition, also known as dysbiosis, contributes to inflammatory disorders^2^. In addition, gut microbiota containment is essential to keep homeostasis and increased microbial translocation is associated with systemic inflammation^3^. The gut barrier may be compromised via physical injury in the mucosa, changes in diet or microbiota composition, host pathology^4^ or as a result of the impairment in the immune barrier^5^. Following gut barrier disruption, commensal or pathogenic bacteria can cross the intestinal epithelium to reach systemic circulation^6^ and tissues such as the liver or gastrointestinal-associated lymphoid tissues among others^7^, favoring the activation of the innate and adaptive immune system^8–11^ Notably, inflammatory bowel disease (IBD), systemic lupus erythematosus (SLE) and cardiovascular diseases have been associated with increased gut permeability^12–15^.

Microbe-associated molecular patterns, microbial products and microbe-derivate metabolites that reach the circulation can modulate the innate immune system^16–19^. Sensing of these microbial signals by myeloid cells may lead to metabolic and epigenetic reprogramming that boosts inflammatory responses to a secondary challenge, a process termed trained immunity (TI)^20,21^. Gut microbiota can induce epigenetic modifications in many host tissues^22,23^. These temporal epigenetic changes can be mediated by microbe-derived metabolites such as short-chain fatty acids (SCFA) through their effects on histone modifying enzymes^22,24,25^. Moreover, metabolic changes produced by alterations of gut microbiota affects the bone marrow (BM) niche^26^. Mice with gut microbiota, compared to Germ-Free (GF) mice, showed increased myelopoiesis^27–29^.

Notably, the long-term effects of TI are explained by the modulation of BMPs^30^. Among the receptors sensing microbial signals, Dectin-1 is a paradigmatic C-type lectin receptor (CLR) that drives TI^31^. Notably, Mincle/*Clec4e^−/-^* is a closely related CLR that shares with Dectin-1 similar signaling pathways^32^ and senses ligands in different microbes^33^, but has not been related to TI. Moreover, the Mincle-Syk axis in dendritic cells (DCs) promotes intestinal barrier integrity upon microbiota sensing^34^.

To learn about the mechanisms connecting gut microbiota translocation with systemic inflammation, we hypothesized that microbiota dissemination upon gut barrier disruption could induce TI in myeloid cells in distant tissues. To investigate this, we administered mice with DSS, which impairs the intestinal barrier, and we found translocation of gut commensals to distal tissues, including the BM. Microbiota translocation drove epigenetic changes and promoted TI in BMPs. *Enterococcus faecalis* (*E. faecalis*) was identified as the predominant species in the BM that activated immune cells in a Mincle/Clec4e-dependent manner. *Clec4e^−/-^* mice showed reduced TI-related inflammation and pathology upon *E. faecalis* administration or DSS treatment. Our results identify the gut microbiota *E. faecalis* as a potential contributor to inflammatory pathologies related to gut barrier disruption through the induction of TI.

## Results

### Gut microbiota translocation induces trained immunity in myeloid bone marrow progenitors

Acute colitis induced by DSS treatment causes massive translocation of gut microbiota to distal tissues^35^. To assess whether gut microbiota translocation could result in enhanced inflammatory response to secondary stimulation, a hallmark of TI, we treated mice for 5 days with 3% DSS in drinking water and, after a 5-day rest, BM cells were collected (Fig. 1A). Bone marrow-derived macrophages (BMDMs) obtained from DSS-treated mice showed a higher capacity to produce TNFα when challenged with either LPS (TLR4 ligand), CpG (TLR9 ligand) or Pam3Csk4 (TLR1/2 ligand) as a secondary challenge compared to non-DSS-treated mice (Fig. 1B). In addition, a higher response against the secondary challenge was found *ex-vivo* in total BM cells stimulated with LPS, CpG or Pam3Csk4 (Fig. S1A). The same result was also observed in BMDMs from DSS mice compared to untreated mice on a *Rag1*^−/-^ background (Fig. S1B) showing that this effect is independent of the adaptive immune system. Administration of antibiotics (ABX) that depleted gut microbiota^36^ prevented the higher TNFα production from DSS mice vs untreated mice upon restimulation of BMDM (Fig. 1C) and total BM (Fig. S1C). To further confirm the role of gut microbiota in BM reprogramming, C57BL/6 Germ-free (GF) mice were administered, or not, a Fecal Microbiota Inoculation (FMI) to restore the conventional gut microbiota before DSS treatment (Fig. 1D). GF mice which received FMI showed increased TNFα production by BMDMs (Fig. 1D) and total BM (Fig. S1D). upon PAMP restimulation.

**Fig.1.**
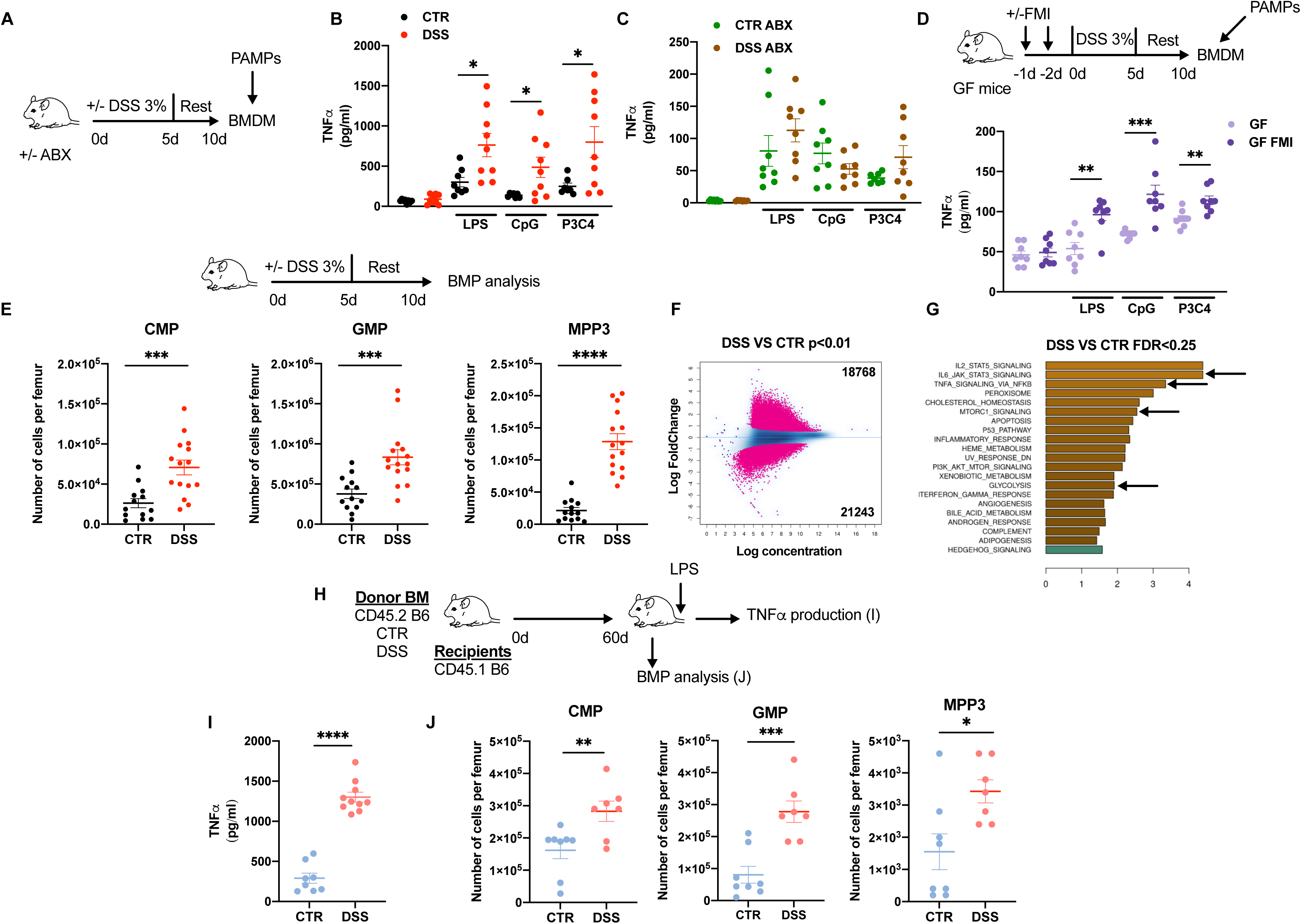
Gut microbiota translocation induces trained immunity in myeloid bone marrow progenitors. (A-C) (A) Mice were treated or not with DSS and antibiotics (ABX) as indicated in the outline, bone marrow (BM) extracted, BM-derived macrophages (BMDMs) generated and stimulated with LPS, CpG, or P3C4, for 24h. (B, C) TNFα production measured by ELISA in culture supernatant of BMDMs from indicated treatments and stimulations. (B) Two pooled independent experiments (n=7-9). (C) Two pooled independent experiments (n=8). (D) Germ-free (GF) mice were inoculated or not with fecal microbiota (FMI) from SPF mice and treated or not with the DSS protocol and BMDM generation and stimulation as in (A). TNFα production measured by ELISA in culture supernatant of BMDMs from indicated treatments and stimulations. Two pooled independent experiments (n=8). (E) Mice were treated or not with DSS as in (A) and bone marrow myeloid progenitors (BMPs) counted at day 10. Graphs show total numbers per femur of common myeloid progenitors (CMP), granulocyte-monocyte progenitors (GMP) and multipotent progenitor cells (MPP3). Three pooled independent experiments (n=13-15). (F,G) ATAC-seq analysis in GMP cells sorted from BM of CTR or DSS mice at day 10 of protocol shown in (A). (F) MAplot showing 40.011 differentially accessible peaks between DSS and CTR mice GMPs, with p<0.01. (G) Bar plot showing enriched gene sets from the MSigDB Hallmark collection with FDR<0.25. Positive normalized enrichment score (NES) values (orange color) and negative (green color) indicate congruent over- or under-expression of a pathway or function-associated genes in DSS compared to CTR (Control) condition. Only the top 20 hits (according to FDR) are represented. A pool of 3 mice was used per biological replicate, and 3 biological replicates were used per condition. (H-J) (H) Donor mice were treated or not with DSS as in (A) and BM grafted to CD45.1 B6 recipient mice. (I) After 60d mice were challenged with LPS and TNFα measured in plasma by ELISA 1h later. Two pooled experiments (n=8-10). (J) Graphs show total numbers per femur of the indicated myeloid BMPs. Two pooled experiments (n=7-8). Individual data and mean ± SEM is shown unless otherwise stated. *p<0.05; **p < 0.01; ***p < 0.005; ****p < 0.001 (Unpaired Student’s t test). See also Figure S1.

Analysis of BMPs showed expansion in common myeloid progenitors (CMP), granulocyte monocyte progenitors (GMP) and multipotent progenitor cells 3 (MPP3), in DSS-treated mice compared with control mice (Fig. 1E; Fig. S1E). To determine if this was due to an intrinsic change in the BMPs by epigenetic reprogramming, the main hallmark of TI, we sorted GMP cells from mice treated or not with the DSS protocol and analyzed chromatin accessibility by ATAC-Seq. Our results revealed deep changes in accessible peaks in GMPs from DSS-treated mice compared to untreated controls (Fig. 1F). Notably, by ascribing open regions to gene promoters, Gene Set Enrichment Analysis (GSEA) showed an enrichment in pathways related to inflammation status such as NF-κB, proinflammatory cytokines, mTOR signaling, and pathways related to glycolysis in DSS-treated compared to untreated mice (Fig. 1G). To confirm that these intrinsic epigenetic changes in BMPs drive TI in the context of DSS treatment and gut microbiota translocation, we assessed whether TI could be transferred by BM graft. BM from donor CD45.2^+^ mice, treated or not with DSS, was grafted into lethally irradiated recipient CD45.1^+^ mice. After 60 days, recipient mice were challenged intraperitoneally (i.p.) with 5μg of LPS to test TNFα production in plasma as a TI readout (Fig. 1H). Mice grafted with BM from DSS-treated donors produced higher amount of TNFα systemically in response to LPS compared to mice grafted with the BM from untreated donors (Fig. 1I). Moreover, myeloid BMPs (CMP, GMP and MPP3) were expanded in mice grafted with BM from DSS-treated donors compared to untreated donors (Fig. 1J). These results show that DSS-driven gut microbiota translocation drives BM epigenetic reprogramming responsible for a long-lasting hyper-responsiveness state and that can be transferred upon BM graft, which defines TI hallmarks.

### Identification of *E. faecalis* as a main driver of bone marrow training upon gut barrier disruption

Following DSS treatment, we found bacterial translocation under anaerobic and aerobic growing conditions not only in the liver, but also in the BM (Fig. S2A,B) This bacterial translocation was abolished in DSS-ABX-treated mice, as expected (Fig. S2A,B). The growth of BM flushed from DSS-treated or control mice in a rich medium for broad-spectrum bacteria revealed one type of morphologically homogenous colony able to grow in aerobic and anaerobic conditions. These homogeneous colonies exhibited positive Gram staining and were unable to grow in MacConkey Agar (medium for selective isolation of Gram-negative and enteric bacilli). Notably, these colonies were positive for the Aesculin test, a selective and differential medium which identifies *Enterococci* (Fig. S2C). This preliminary screening identified the bacteria present in DSS-treated mice BM as a Gram-positive species from the genus *Enterococcus* (Fig. S2C). To confirm the genus, an API strep test was carried out, concluding that all colonies detected in the BM belonged to the same genus, *Enterococcus sp*., and identified the species with 99.3% certainty as *E. faecalis* (Fig. 2A). To confirm the species, MALDI-TOF mass spectrometry analysis was performed, concluding that the translocated bacteria was *E. faecalis* (Fig. 2B). The quantification of *E.faecalis* translocated to the BM of one tibia and femur revealed an average growth of 2×10^2^ CFU both in aerobic and anaerobic conditions (Fig S2D).

**Fig.2.**
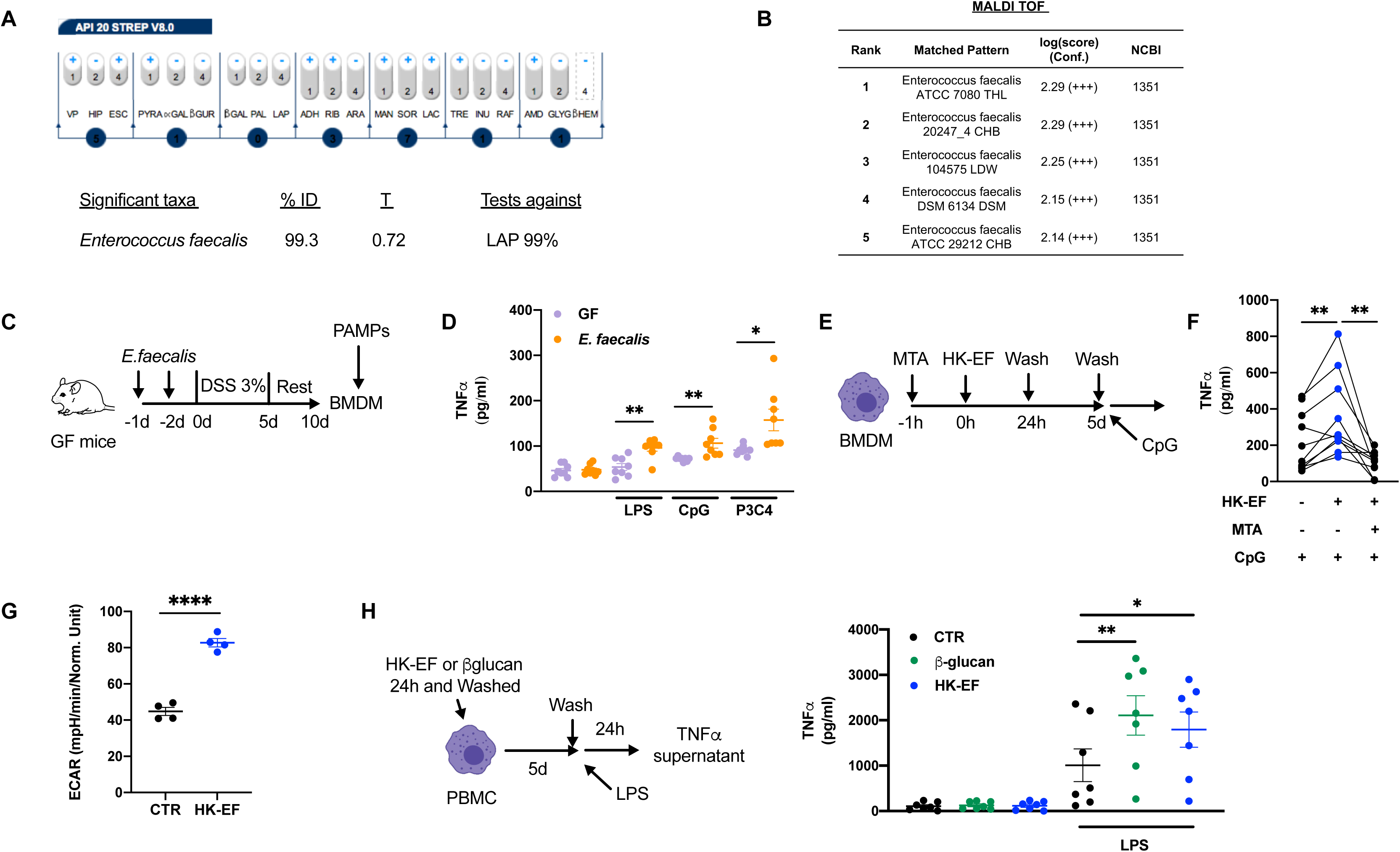
Identification of Enterococcus faecalis as a main driver of bone marrow training upon gut barrier disruption. (A, B) BM from DSS-treated mice was flushed and cultured in LB plates. (A) Result of a representative API 20 strep test read of growing bacterial colonies after 4h (VP to LAP tests), and 24h (ADH to GLYG tests) of bacterial culture. The result is indicated as positive or negative based on colorimetric changes. (B) MALDI biotyper report showing numerical(B) Numerical log score sidentifying species of bacteria detected by MALDI-TOF analysis of bacteria colonies obtained from LB plates in aerobic conditions. A score of ≥1.80 represents high confidence and it is accepted for bacterial identification. Was (C-D) (C) GF mice were monocolonized or not with *E. faecalis,* and treated with DSS, BM extracted, BMDMs generated and stimulated with LPS, CpG or P3C4 for 24h. (D) TNFα production measured by ELISA in culture supernatant of BMDMs from indicated treatments and stimulations. Two pooled experiments (n=8). (E-F) (E) BMDMs obtained from C57BL/6J mice, were pretreated or not with the methyltransferase inhibitor, 5′-methylthioadenosine (MTA) for 1h before incubation with heat killed *E. faecalis* (HK-EF) for a further 24h. After this time, media was changed, and cells were rested for 5 days. (F) TNFα production measured by ELISA in culture supernatant of BMDMs after 24h stimulation with CpG. Three pooled experiments (n=10). (G) Extracellular Acidification Rate (ECAR) in BMDMs after 24 hours stimulation with or without HK-EF measured by performing a MitoStressTest. n=4. (H) Human peripheral blood mononuclear cells (PBMC) were incubated 24h with or without of β-glucan or HK-EF, washed and left resting for 5 days before restimulation with LPS (left) and TNFα concentration was measured in the supernatant of human PBMC upon LPS restimulation (right). Two pooled experiments (n=7). Individual data and mean ± SEM is shown unless otherwise stated. Unpaired Student’s t test was used to compared conditions in (D) and (H). Paired Student’s t test was used to compare conditions in (F). *p < 0.05; **p < 0.01; ****p < 0.001. See also Figure S2.

Next, we tested the capacity of *E. faecalis* translocation *per se* to induce TI. To this aim, GF mice were monocolonized with 10^8^ CFU *E. faecalis*, for two consecutive days prior to DSS administration (Fig. 2C). Compared with GF mice, the monocolonization with *E. faecalis* was sufficient to induce enhanced TNFα production by BMDMs upon rechallenge with LPS, CpG and Pam3Csk4 (Fig. 2D).

We next assessed the ability of heat-killed *E. faecalis* (HK-EF) to induce TI hallmarks *in vitro*. Following the typical *in vitro* TI protocol^37^, we tested *E. faecalis* as training stimulus in BMDMs, priming for 24h with HK-EF, washing, resting for 5 days and re-stimulating with CpG. Preincubation with 5′-methylthioadenosine (MTA), a methyltransferase-selective inhibitor, was used to inhibit TI-related epigenetic modifications (Fig. 2E). We found that priming of BMDM with HK-EF increased TNFα production after the re-challenge with CpG, and the effect was abolished in presence of MTA (Fig. 2F). Moreover, BMDMs stimulated with HK-EF for 24 hours showed increased extracellular acidification rate (ECAR), indicative of glycolytic activity, compared to unstimulated BMDMs, another hallmark of TI^38,39^ (Fig. 2G). Notably, these results were reproduced in peripheral blood mononuclear cells (PBMC) from human buffy coats, where HK-EF*-*primed cells produced more TNFα after re-challenge with LPS, compared to non-trained cells and to a similar extent than β-glucan training (Fig. 2H). We conclude that, upon intestinal barrier disruption, *E. faecalis* translocate to the BM, where they induce trained immunity in myeloid cells and their precursors.

### Systemic administration of heat-killed *E. faecalis* trains myeloid progenitors

We then evaluated the ability of HK-EF to induce TI *in vivo* in the absence of intestinal barrier disruption. C57BL/6 male mice were injected intravenously (i.v.) with a range of HK-EF doses. As a readout for TI *in vivo*, mice were challenged i.p. with LPS and TNFα measured in plasma (Fig. 3A). Notably, the i.v. injection of HK-EF resulted in a bell-shaped curve for induction of TNFα production upon restimulation with LPS, being the optimal dose 2×10^3^ CFU (Fig. S3A). In addition, we found that the duration of the priming effect was at least 8 weeks (Fig. S3B, C).

**Fig 3.**
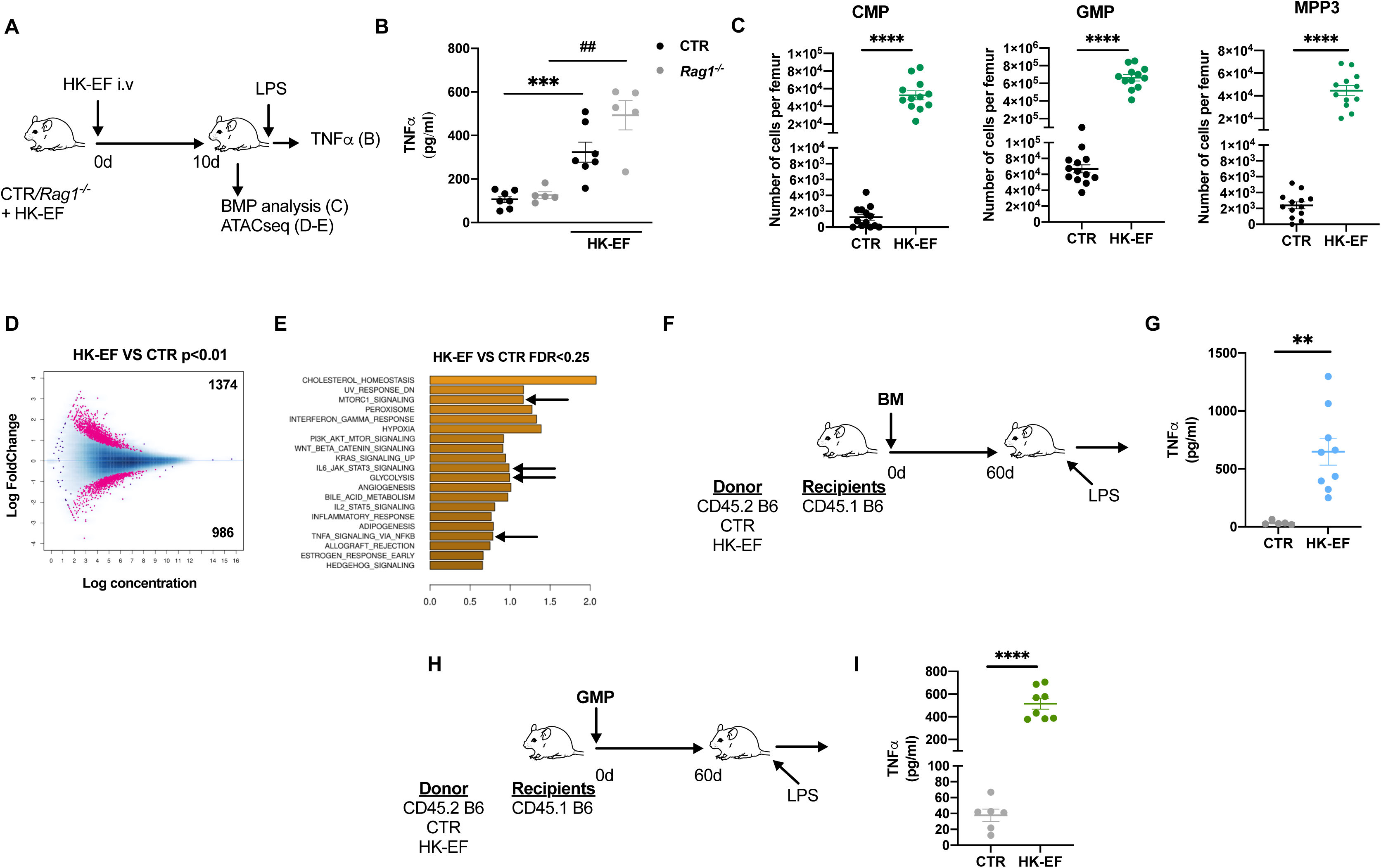
Systemic administration of heat-killed *E. faecalis* trains myeloid progenitors. (A, B) (A) Wild type (WT) or *Rag1^−/-^* mice were treated or not with HK *E. faecalis* (HK-EF) i.v. followed by a challenge with LPS 10d later. (B) TNFα concentration measured by ELISA in plasma obtained 1h after LPS stimulation in the indicated genotypes and treatments. Two pooled independent experiments (n=5-7). (C) Mice were treated or not with HK-EF i.v. and BMPs analyzed 10d later. Graphs show total cell numbers per femur of common myeloid progenitors (CMP), granulocyte-monocyte progenitors (GMP) and multipotent progenitor cells (MPP3). Three pooled independent experiments (n=12-13). (D-E) ATAC-seq analysis in GMP cells sorted from BM of untreated (CTR) or HK-EF treated mice at day 10 of protocol shown in (A). (D) MAplot showing 2.360 differentially accessible peaks between HK-EF and CTR mice GMPs, with p<0.01. (E) Bar plot showing enriched gene sets from the MSigDB Hallmark collection with FDR<0.25. Positive normalized enrichment score (NES) values (orange color) and negative (green color) indicate congruent over- or under-expression of a pathway or function-associated genes in HK-EF compared to CTR condition. Only the top 20 hits (according to FDR) are represented. A pool of 3 mice was used per biological replicate, and 3 biological replicates were used per condition. (F-G) (F) Donor mice were treated or not with HK-EF and rested for 10d as in (A) and BM grafted to CD45.1 B6 recipient mice. After 60d mice were challenged with LPS (G) TNFα concentration measured by ELISA in plasma obtained 1h after LPS stimulation. One representative experiment of two performed (n=5-9). (H) Donor mice were treated or not with HK-EF as in (C) and GMP grafted to CD45.1 B6 recipient mice. After 60d mice were challenged with LPS. (I) TNFα concentration measured by ELISA in plasma obtained 1h after LPS stimulation. Two pooled independent experiments (n=6-8). Individual data and mean ± SEM from indicated independent experiments is shown unless otherwise stated. Unpaired Student’s t test was used to compare groups, unless otherwise stated *p < 0.05; ***p < 0.005; ****p < 0.001; ^##^p < 0.01 vs *Rag1^−/-^*. See also Figure S3.

Using the lowest effective dose of HK-EF that induced TNFα production upon restimulation with LPS (Fig. S3A) we also found this TI readout in a *Rag1^−/-^* background (Fig. 3B), showing that this effect is independent of B and T cells. The i.v. injection of HK-EF also resulted in expansion of BMPs (Fig. 3C). To determine if this was due to an intrinsic change in epigenetic reprogramming, we sorted GMP cells from mice treated with the HK-EF or control mice and analyzed chromatin accessibility by ATAC-Seq. Our results revealed differentially accessible peaks in GMPs from HK-EF treated mice compared to control mice (Fig. 3D). By ascribing open regions to gene promoters, GSEA showed an enrichment in pathways related to inflammation status such as proinflammatory cytokines, mTOR and pathways related to glycolysis in HK-EF compared to untreated mice (Fig. 3E). To test the role of the intrinsic epigenetics changes in BM myeloid progenitors induced by HK-EF, we performed a BM graft using donors which had been i.v. injected with HK-EF 10 days before vs untreated control donor mice (Fig. 3F). After 60 days, recipient mice were challenged i.p. with 5μg of LPS to test TNFα production in plasma as a TI readout. Mice grafted with BM from donor primed with HK-EF showed a boost in the secondary response *in vivo* upon LPS rechallenge in the recipient mice, compared to mice grafted with BM donated by control mice (Fig. 3G). Of note, a similar experiment reconstituting lethally-irradiated recipient mice exclusively with sorted GMP cells from donor mice treated or not with HK-EF (Fig. 3H) showed an enhanced TNFα production after the LPS challenge when donor GMP cells came from treated mice (Fig. 3I), showing the key role of epigenetics changes found in GMP. In conclusion, i.v. administration of HK-EF induces long-term trained immunity that can be transferred by BM progenitors.

### Heat-killed *E. faecalis* triggers Mincle-dependent trained immunity in BMDMs

Since Dectin-1 is a paradigmatic C-type lectin receptor (CLR) that induces TI^40^ and CLRs can bind glycolipids in bacteria^34^ we analyzed the effect of inhibitors related to the CLR signaling pathway in HK-EF-induced TI. BMDMs from WT mice were pre-treated with inhibitors for one hour prior to priming with HK-EF and the effect in enhanced TNFα production upon restimulation with CpG 5d later was evaluated (Fig. 4A). Notably, inhibition of either Syk (Fig. 4B), CARD9 (Fig. 4C) or mTOR (Fig. 4D) abolished the enhanced secondary response induced by priming of BMDMs with HK-EF *in vitro*. Since several CLRs are upstream this signaling pathway^32^, we used a NFAT-reporter system^41,42^ to determine which CLRs can detect ligands in HK-EF. B3Z cells expressing the extracellular domains of either Dectin-1, Dectin-2 or Mincle coupled to an intracellular CD3ζ domain to activate the NFAT reporter were exposed to known ligands for each receptor, as positive controls, and HK-EF. Notably, while the reporter signal was triggered by specific ligands for all receptors, only Mincle reporter was activated by HK-EF (Fig. 4E). Moreover, a limiting dose of HK-EF not only triggered the reporter activity through the Mincle-CD3ζ chimera, but also through the natural Mincle receptor in a B3Z cell line transfected with FcRγ and Syk^42^, indicating that HK-EF triggers the Mincle/Syk pathway (Fig 4F). As an alternative approach to validate this result, we stained HK-EF with Mincle-ectodomain-human-Fc chimera (Mincle-hFc), which bound specifically to a fraction of the bacteria exposing the ligand, compared with the control human-Fc chimera (Fig. 4G). In addition, BMDMs treated with HK-EF showed increased S6 ribosomal protein phosphorylation (pS6) indicative of mTOR activation, which was dependent on Mincle since phosphorylation was abolished in BMDM *Clec4e-/-* (Fig 4H). To test the role of Mincle in the induction of TI by HK-EF, we generated BMDMs from WT and *Clec4e*^−/-^ mice. Following priming with HK-EF, the enhanced TNFα production upon restimulation with CpG 5d later observed in WT was prevented in BMDM from *Clec4e*^−/-^ mice (Fig 4I). We conclude that HK-EF induces TI through the Mincle-Syk pathway that results in mTOR activation, a signaling pathway linked to increased TI^38^.

**Fig. 4.**
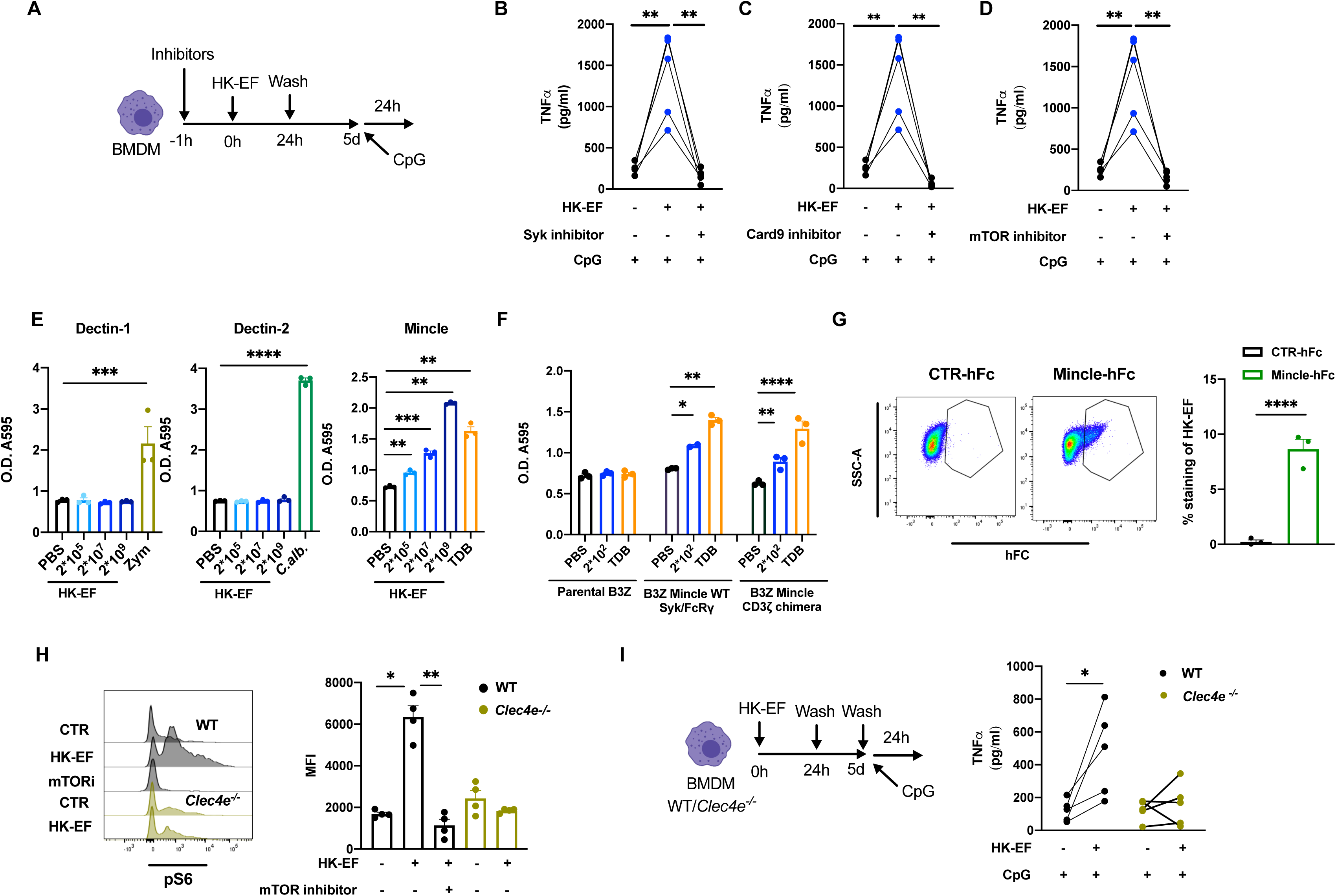
Heat-killed *E. faecalis* triggers Mincle-dependent trained immunity in BMDMs. (A-D) (A) BMDMs were treated or not with HK-EF upon pretreatment with SYK inhibitor (B), Card9 inhibitor (C) or mTOR inhibitor (D) and stimulated with CpG 5d later, TNFα production measured by ELISA in culture supernatant of BMDMs 24h after CpG stimulation. Paired Student’s t test comparing independent BMDM cultures (n=5). (E) NFAT reporter activity in response to the indicated stimuli and increasing doses of HK-EF in B3Z cells expressing the extracellular domains of either mouse Dectin-1 (left), Dectin-2 (middle), or Mincle (right) coupled to an intracellular CD3ζ. Combined results from 3 independent experiments are shown. Statistical analysis was performed using one-way ANOVA with Bonferroni post-hoc test. (F) NFAT reporter activity in response to TDB or the indicated dose of HK-EF in parental B3Z cells or, mouse WT Mincle receptor in B3Z co-expressing Syk and FcRγ or B3Z expressing human Mincle-CD3ζ chimera. Results shown are a combination of 3 independent experiments. Statistical analysis was performed by one-way ANOVA with Bonferroni post-hoc test. (G) Representative plots (left) and graph depicting the frequency (right) of HK-EF stained with control-hFc or Mincle-hFc. Pool of three independent experiments. (H) Representative histograms (left) and MFI (right) of phosphorylation of S6 ribosomal protein (Ser235/236) measured by flow cytometry in BMDMs from WT and *Clec4e^−/-^* mice stimulated or none 24h with HK-EF. Pool of two independent experiments. (I) BMDMs from WT or *Clec4e^−/-^*mice were treated with HK-EF or PBS according to graphical outline. TNFα production measured by ELISA in culture supernatant of BMDMs from indicated treatments and stimulations. Paired Student’s t test compares 5 independent WT and *Clec4e ^−/-^* BMDM cultures. Individual data and mean ± SEM from indicated independent experiments is shown unless otherwise stated. * p < 0.05; ** p < 0.01; *** p < 0.005; **** p < 0.001.

### Mincle receptor stimulation induces trained immunity

Despite the similarities of Dectin-1 and Mincle signaling pathways, Mincle has not been previously linked to the induction of TI. We thus wondered whether a well-established Mincle agonist such as Trehalose-6,6-dibehenate (TDB)^43^ could also induce TI. Following BMDM treatment with TDB for 24h, cells were rested for 5d and re-challenged with CpG (TLR9 agonist) (Fig. 5A). Pre-treatment with TDB enhanced TNFα secretion by BMDMs, an effect that was prevented by co-incubation with the epigenetic inhibitor MTA during the pre-treatment (Fig. 5B). The enhanced secondary response against CpG induced by TDB pretreatment was abolished in BMDM from *Clec4e^−/-^* mice (Fig. 5C). Moreover, 24h of TDB stimulation produced an increase in basal glycolysis in BMDMs (Fig. 5D). Notably, BMDMs treated with TDB exhibited increased pS6 in a Mincle-dependent fashion (Fig. 5E), further confirming the key role of Mincle receptor in the activation of mTOR pathway in BMDMs, a signaling pathway that contributes to TI^38^. To analyze the ability of TDB to induce TI *in vivo* through Mincle, WT and *Clec4e^−/-^* mice were i.v. injected with dimethyl dioctadecyl ammonium bromide (DDA) liposomes containing or not TDB (DDA vs DDA:TDB) to induce TI and re-challenged i.p. with LPS 10d later (Fig. 5F). Mice pre-treated with DDA:TDB showed increased production of TNFα compared with DDA pretreatment alone in WT, showing TI induction by DDA:TDB that was prevented in *Clec4e^−/-^* mice (Fig. 5G). We conclude that the activation of Mincle by selective agonist TDB meets signaling, metabolic and functional hallmarks of TI in myeloid cells, affecting BMPs,

**Fig 5.**
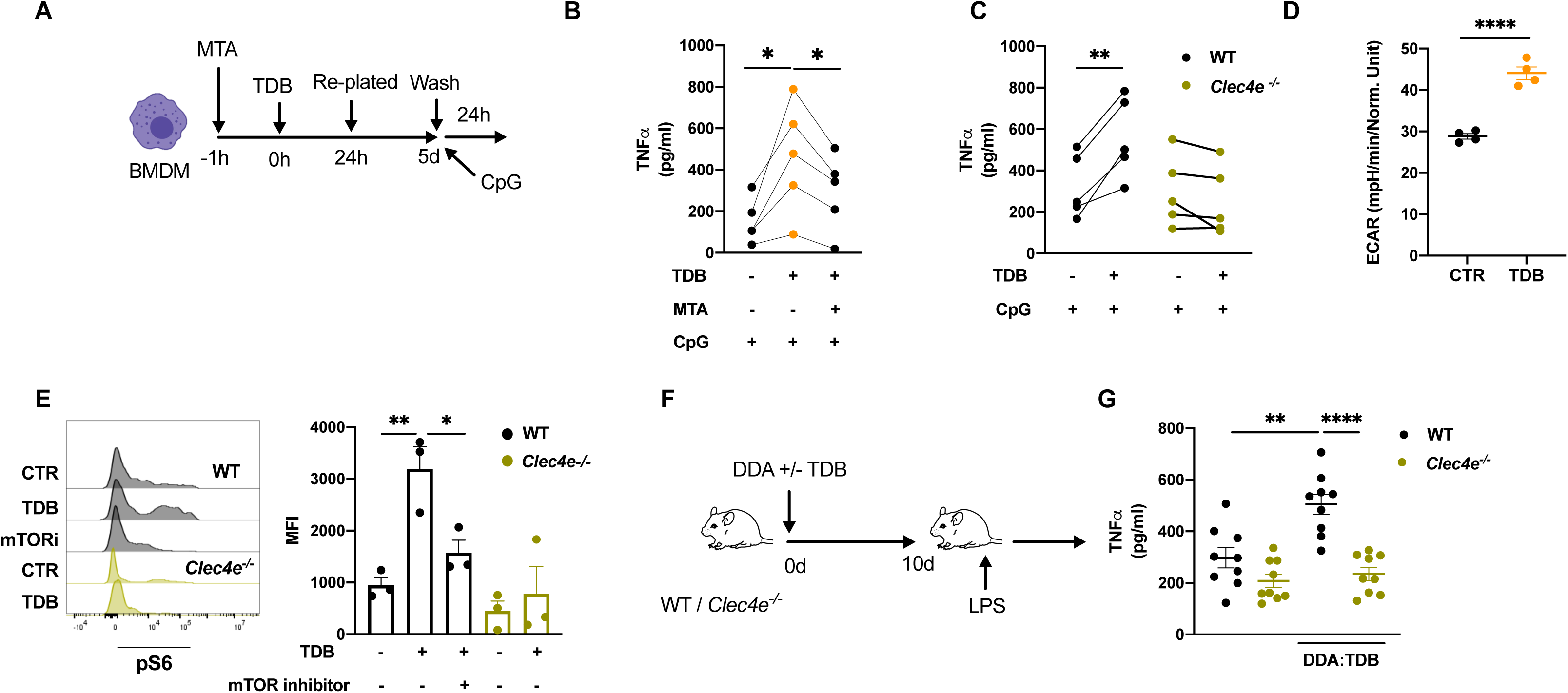
Mincle receptor stimulation induces trained immunity. (A, B) (A) BMDMs were treated or not with MTA 1h before plating in Trehalose-6,6-dibehenate (TDB)-coated wells. After 24 hours cells were re-plated in non-TDB coated plates and rested for 5 days before challenge with CpG for 24 hours. (B) TNFα production measured by ELISA in culture supernatant of BMDMs from indicated treatments and stimulations. Paired Student’s t test compares 5 independent cultures. (C) BMDMs from WT or *Clec4e^−/-^* mice were treated as described in (A). TNFα production measured by ELISA in culture supernatant of BMDMs from indicated treatments and stimulations. Paired Student’s t test compares 5 independent WT and *Clec4e ^−/-^*BMDM cultures. (D) Extracellular Acidification Rate (ECAR) of BMDMs stimulated or not with TDB for 24 hours measured in the basal step of MitoStress Test in a Seahorse assay (n=4). (E) Representative histograms (left) and MFI (right) of phosphorylation of S6 ribosomal protein (Ser235/236) measured by flow cytometry in BMDMs from WT and *Clec4e^−/-^* mice stimulated or not for 24h with TDB. Pool of 3 independent experiments. (F-G) (F) WT or *Clec4e^−/-^* mice were treated with DDA or DDA:TDB i.v. and restimulated with LPS 10d later. (G) TNFα concentration measured by ELISA in plasma obtained 1h after LPS stimulation. Three pooled independent experiments (n=9). Individual data and mean ± SEM is shown unless otherwise stated. Unpaired Student’s t test was used to compare conditions unless otherwise stated. *p<0.05; **p<0.01; *** p < 0.005; ****p < 0.001.

### Heat-killed *E. faecalis* induces Mincle-dependent trained immunity *in vivo* that is protective against fungal infection

To address the role of Mincle receptor in HK-EF-mediated TI *in vivo*, WT and *Clec4e^−/-^*mice were primed with HK-EF and 10d later challenged with LPS to test the secondary response (Fig. 6A). WT mice pretreated with HK-EF exhibited an enhanced TNFα production following LPS rechallenge that was fully abolished in *Clec4e*^−/-^ mice (Fig. 6B). Moreover, priming with HK-EF resulted in a Mincle-dependent expansion of BM progenitors (Fig. 6C).

**Fig 6.**
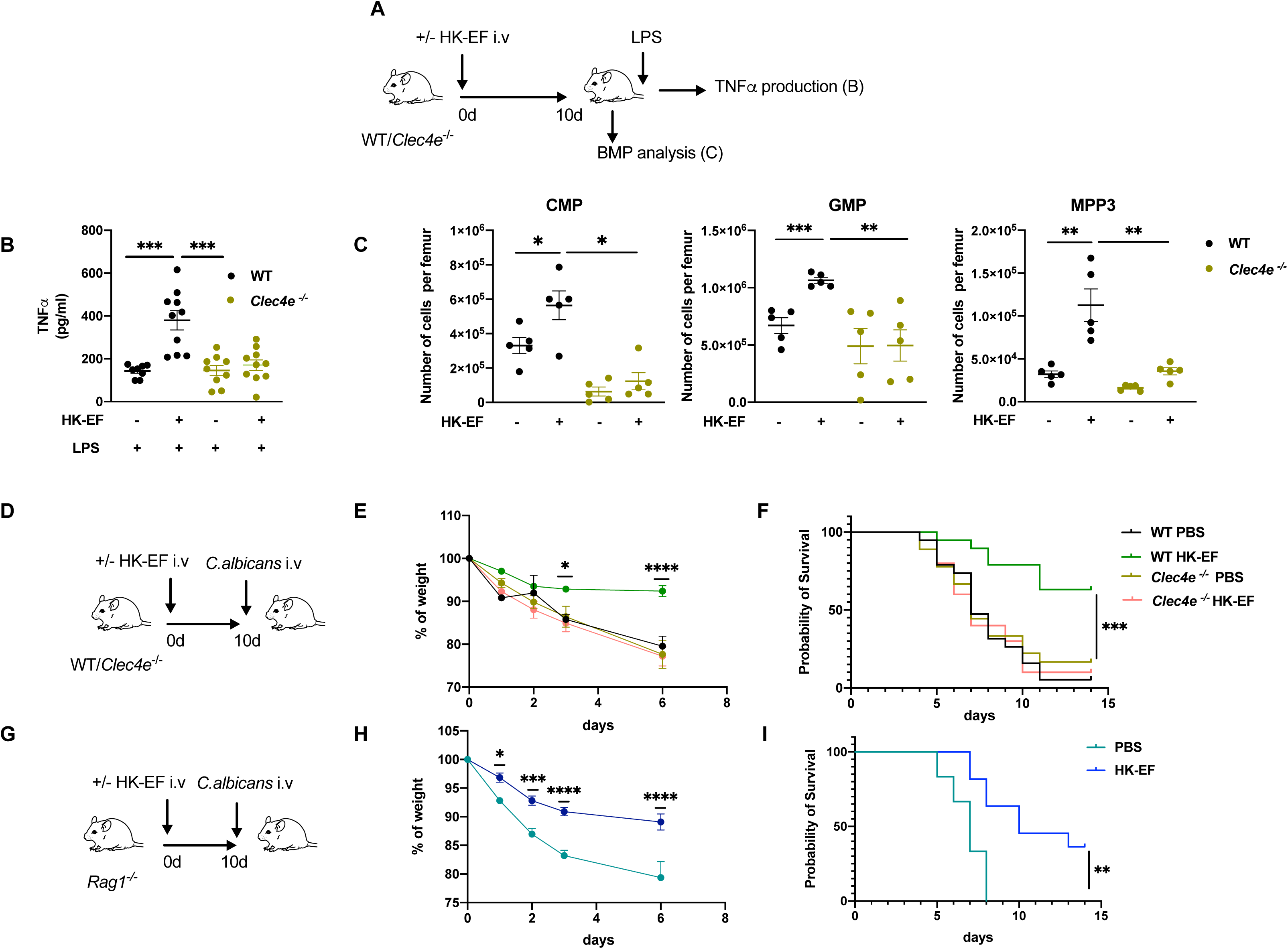
Heat-killed *E. faecalis* induces Mincle-dependent trained immunity *in vivo* that is protective against fungal infection. (A, B) (A) WT and *Clec4e ^−/-^* mice were treated or not with HK-EF i.v. and challenged with LPS i.p at day 10. (B) TNFα concentration in plasma from mice of the indicated treatments and genotypes. Three pooled independent experiments (n=8-10). (C) WT and *Clec4e ^−/-^* mice were treated or not with HK-EF i.v. and BMPs counted at day 10..Graphs show total numbers per femur of common myeloid progenitors (CMP), granulocyte-monocyte progenitors (GMP) and multipotent progenitor cells (MPP3). One representative experiment (n=5). (A-C) Individual data and mean ± SEM is shown. Unpaired Student’s t test was used to compare conditions. *p<0.05; **p<0.01; *** p < 0.005. (D-F) WT and *Clec4e ^−/-^* mice treated or not with HK-EF were challenged with *C. albicans* at day 10. (E) Weight loss and (F) survival curves are shown. (G-I) WT and *Rag1^−/-^* mice treated or not with HK-EF were challenged with *C. albicans* at day 10. (H) Weight loss and (I) survival curves are shown. (D-I) Results from a pool of two independent experiments ((n = 10/group). Two-way ANOVA test comparing CTR vs HK-EF weight. Log rank (Mantel-Cox) test for survival curve comparison. *p<0.05; **p<0.01; *** p < 0.005; ****p < 0.001. See also Figure S4.

Next, we explored the potential effect of TI induced by HK-EF via Mincle in prevention of fungal infection. WT and *Clec4e*^−/-^ mice were i.v. trained or not with HK-EF (2×10^3^ CFU) and after 10d mice were infected i.v. with a lethal dose of *Candida albicans* (Fig. 6D). Training with HK-EF resulted in reduced weight loss and improved survival compared to non-trained mice, a protective effect that was dependent on Mincle (Fig. 6E, F). Notably, this protective effect was independent of T and B cells (Fig. 6G), since HK-EF-trained *Rag1^−/-^* mice also showed reduced morbidity and mortality compared with non-trained *Rag1^−/-^* mice (Fig. 6H, I). Moreover, we tested whether intranasal (i.n) administration of HK-EF, similar to other bacteria-based preparations that induce TI such as MV130 and MV140^39,44,45^, could be protective. WT mice were treated or not with HK-EF (2×10^6^ CFU) i.n, three times per week for two consecutive weeks, as described previously for i.n. polybacterial vaccines^39,45^ (Fig. S4A). Seven days after the last dose mice were i.v. infected with a lethal dose of *C. albicans.* We found that i.n. HK-EF pretreatment reduced the mortality compared with untreated mice (Fig.S4A). Our results show that HK-EF induces TI *in vivo* via Mincle, resulting in protective immunity against heterologous infections, regardless of administration route.

### Mincle deficiency prevents TI and pathology associated to DSS-induced gut barrier disruption

We thus wondered whether TI induced by gut bacterial translocation would also depend on Mincle expression. Following DSS treatment to increase gut permeability in WT and *Clec4e*^−/-^ mice, we generated BMDMs and treated them with LPS, CpG or P3C4 (Fig. 7A). In agreement with our results above (Fig. 1B) this stimulation resulted in boosted TNFα production in BMDMs obtained from DSS-treated compared to untreated mice, a boosting effect that was dependent on Mincle (Fig. 7B). Moreover, the expansion of CMP, GMP and MPP3 found in WT mice upon DSS treatment was prevented in *Clec4e*^−/-^ mice (Fig. 7C).

**Fig. 7.**
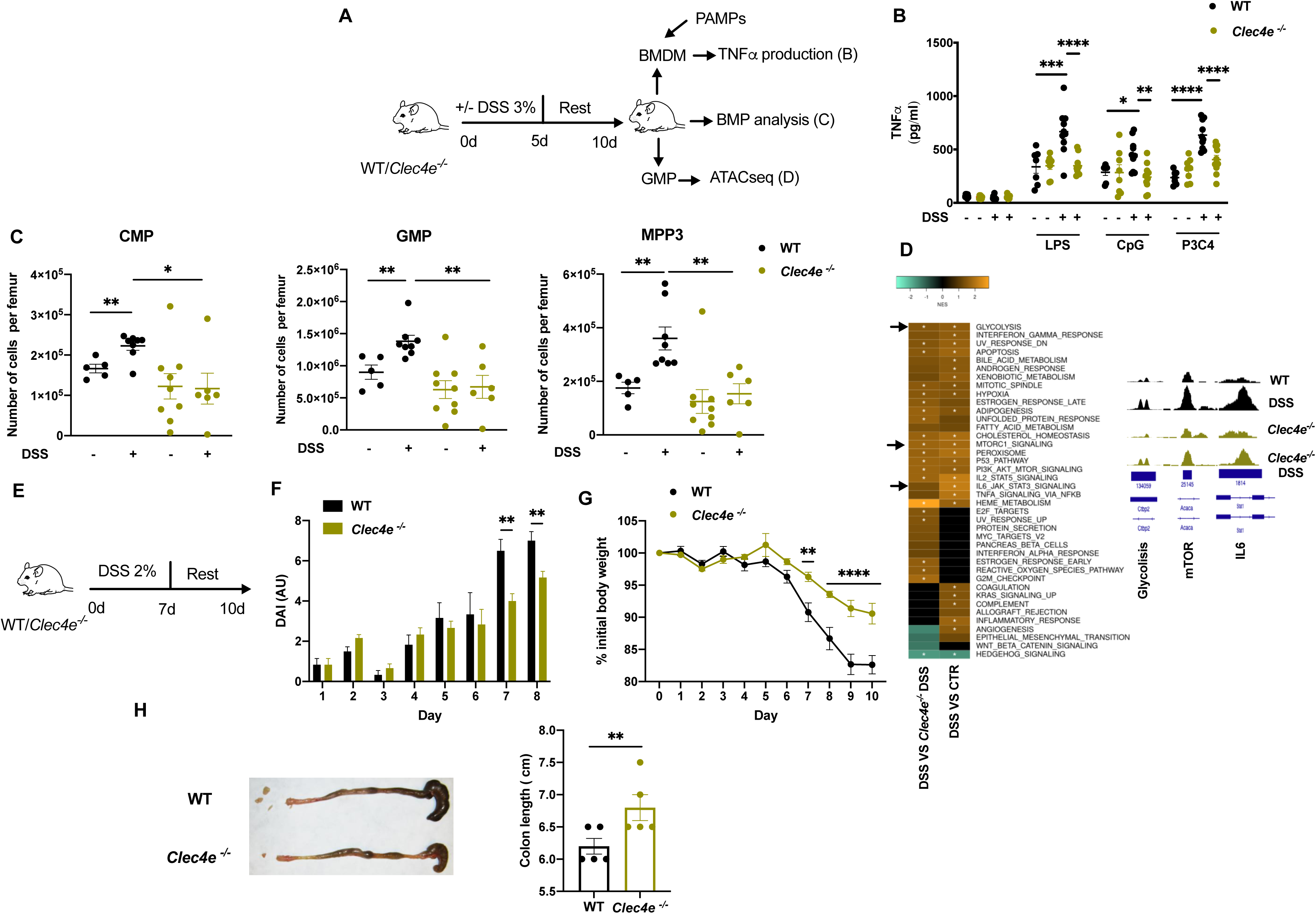
Mincle deficiency prevents trained immunity and pathology associated to DSS-induced gut barrier disruption. (A, B) (A) WT and *Clec4e ^−/-^* mice were treated or not with DSS for 5d and rested. At day 10 BM was extracted, BMDMs generated and stimulated with LPS, CpG, or P3C4 for 24h. (B) TNFα production measured by ELISA in culture supernatant of BMDMs from indicated treatments and stimulations. Two pooled independent experiments (n=7-10). (C) Mice were treated or not with DSS as in (A) and BMPs counted at day 10. Graphs show total numbers per femur of common myeloid progenitors (CMP), granulocyte-monocyte progenitors (GMP) and multipotent progenitor cells (MPP3). Two pooled independent experiments (n=5-9). (D) Mice were treated or not with DSS as in (A) and GMPs were sorted from BM at day 10 of protocol shown in (A) and analyzed by ATAC-Seq. Left: Comparative heatmap representing normalized enrichment score (NES) values for gene sets from the MSigDB Hallmark collection, as calculated by GSEA, after ascribing open regions to gene promoters. Orange and green colors represent significant enrichment with FDR < 0.25. White asterisks indicate FDR < 0.05. Positive NES values (orange color) and negative (green color) indicate congruent change of pathway or function-associated genes in DSS vs CTR, or DSS vs *Clec4e^−/-^*DSS, as indicated. Right: representative open regions modulated in pathways associated to TI. A pool of 3 mice was used per biological replicate, and 3 biological replicates were used per condition. (E-H) (E-H) (E) WT and *Clec4e ^−/-^* mice were treated with DSS as indicated.(F) Disease activity index (DAI) (n=8). (G)weight loss, (n=10) and (H) Colon length (n=5) in WT and *Clec4e ^−/-^* mice. Two pooled independent experiments. Individual data and mean ± SEM is shown unless otherwise stated.. Unpaired Student’s t test was used to compare groups; *p<0.05; **p<0.01; *** p < 0.005; ****p < 0.001.

To determine if DSS-induced changes in epigenetic reprogramming (Fig. 1F, G) were dependent on Mincle, we sorted GMP cells from WT or *Clec4e^−/-^* mice treated or not with the DSS protocol and analyzed chromatin accessibility by ATAC-Seq. By ascribing open regions to gene promoters, GSEA showed an enrichment in pathways related to inflammation status, such as proinflammatory cytokines, mTOR and pathways related to glycolysis in DSS-treated compared to untreated mice (Fig. 7D). Notably, these pathways were also enriched in DSS-treated WT mice compared with DSS-treated *Clec4e^−/-^*mice (Fig. 7D), indicating that DSS-induced epigenetic reprogramming is dependent on Mincle expression.

A pathological outcome of DSS treatment is the induction of colitis. We thus tested if the absence of Mincle could affect colitis development following a standard DSS-colitis protocol^46^ (Fig. 7E). *Clec4e*^−/-^ mice showed reduced disease activity index, weight loss and colon inflammation upon DSS-induced colitis compared with WT (Fig. 7F-H). These results show that Mincle absence improves IBD outcome, providing a potential target to treat pathologies associated with disrupted gut barrier integrity.

## Discussion

Increased gut permeability leading to bacterial translocation can arise after exposure to certain diet components, medications, alcohol, radiation, hyperglycemia or as a consequence of pathologies such as type 2-diabetes mellitus, autoimmune disease (IBD or SLE), or ischemia after stroke, among other factors^12–15^. A functional outcome of gut microbiota dissemination is a systemic immune response with the production of systemic IgG specific for the translocated microbiota, as well as to mucosal IgA^47^. In addition, western diet-driven alterations in gut microbiota are linked to increased gut barrier permeability^48–49^. Notably, western diet feeding triggers systemic inflammation and myeloid cell proliferation and epigenetic reprogramming, namely TI^50^. Herein, we hypothesized that gut microbiota translocation could induce TI in BMPs. We showed that a hard disruption in intestinal barrier integrity allowed bacterial translocation to the BM, inducing a TI phenotype characterized by epigenetic rewiring in GMPs, with a concomitant expansion of myeloid progenitor cells. Under IBD-like conditions, activation of the adaptive immune system drives local and systemic inflammation, generating high concentration of proinflammatory cytokines^51,52^. Notably, the enhanced inflammatory response to secondary challenges was maintained in mice lacking B and T cells (*Rag1^−/-^* mice), pointing towards a phenotype driven by innate immune cells. TI is a long-lasting process, intrinsic to BM cells are therefore transferable by bone marrow graft^53,54^. Accordingly, we found that BM progenitor cells grafted from DSS-treated mice to healthy mice transfer the trained phenotype, inducing higher cytokine production in response to secondary challenge with LPS up to 60 days after the graft, showing that bacterial translocation induces long-term and transferable changes in BM progenitor cells.

We could purify just one translocated species in the BM of DSS-treated mice, identified as *E. faecalis*, which is an opportunistic bacteria, able to induce infection under weakened immune system conditions^55^. *E. faecalis* is more abundant in the gut microbiota from IBD patients compared to healthy patients, but its potential role in the IBD pathology remains unknown^56^. Of note, translocation of *E. faecalis* is associated to gastric acid suppression in alcoholic liver disease, leading to liver inflammation and hepatocyte death^57^. Moreover, the translocation of other species of *Enterococcus*, such as *E. gallinarum*, has been associated with antibody response to a specific subset of lupus autoantigens^8^. We describe here that *E. faecalis* translocation trained BMPs and enhanced their secondary response against TLR agonist. *In vitro*, HK-EF primed hyperresponsiveness and metabolic reprogramming of myeloid cells, characterized by induction of glycolytic metabolism, as described for other TI inductors such as β-glucan^37,58^, BCG^59^ or MV130^39^. Besides, HK-EF trained human PBMC as described for others TI-inducing stimuli such as MV130^39^ or β-glucan ^60,37^. We show that HK-EF induced TI *in vivo*, characterized by an expansion in myeloid progenitor cells, epigenetic changes in GMP, enhanced response upon restimulation and independent of an adaptive immune response. The TI phenotype persists and can be transferred by BM transplantation from trained mice to non-trained recipient mice^61^. Indeed, HK-EF induced long-lasting epigenetic changes in GMP. Similar to DSS treatment, we found that the TI phenotype was transferred by graft of GMP from HK-EF primed mice to non-trained recipient mice, with enhanced secondary responses detected up to 60d after the graft.

We describe that a pathway involving Syk, Card9 and mTOR was essential for the TI effect produced by HK-EF. We identify Mincle as the upstream receptor coupled to this signaling pathway, specifically sensing *E. faecalis* and responsible for TI induction. Mincle is a well-characterized sensor for a diversity of microbial ligands that can mediate inflammatory responses^43,62–64^. Mincle can also detect microbiota signals and regulate intestinal immunity and mucosal barrier function^34^. Moreover, Mincle signaling promotes intestinal inflammation^65^. One of the main pathologies associated with bacterial translocation is IBD^66,67^. The two main clinical manifestations of IBD are Crohn’s disease and ulcerative colitis^68^. Biopsies from IBD patients, DSS-induced mouse colitis model and TNBS-induced mouse colitis model show increased expression of Mincle, correlating with gut inflammation. Deficiency of Mincle attenuates experimental colitis, while administration of the Mincle agonist TDB exacerbated the disease^65^. Our results showing that TDB and HK-EF, through Mincle, induced TI hallmarks suggest the possibility that TI may contribute to gut inflammation. Combinative approaches targeted to CLRs and other PRRs have been suggested for fighting IBD^69^. Our results suggest that the blockade of TI induced by Mincle activation due to bacterial translocation is a potential target to improve TI-related diseases.

In addition, TI has been targeted as a potential immunotherapy tool to improve protection against heterologous infections^45,70–72^. We found that i.v. or i.n administration of HK-EF protected against *Candida albicans* infection. This effect is Mincle dependent and independent of the adaptive immune system. Thus, HK-EF could be used as a Trained Immunity-based Vaccine (TIbV) to prevent heterologous infections, similar to polybacterial preparations MV130^39,73,74^ and MV140^44,75^, or BCG^76^. The protective effect of low doses of HK-EF through different administration routes (i.v. and i.n.) suggests its use in immunotherapy against infectious diseases.

Although we have found that HK-EF trains human PBMCs ex vivo, some **limitations of our study** are related to the impossibility to obtain BMPs from IBD patients to evaluate whether these patients show TI hallmarks in their BMPs. Similarly, we have not tested the capacity of HK-EF to induce protection against infection in humans, which would require a clinical assay. Of note, MV140, a polybacterial preparation that contains *E. faecalis* in its formulation, is protective against recurrent urinary tract infections in clinical assays^77–79,80^. An additional limitation is that while *Clec4e^−/-^* mice indicate a role for Mincle, the deletion is not cell-specific and the critical cell type where Mincle is sensing HK-EF is not fully identified. Nevertheless, the expression of Mincle receptor is quite restricted to some myeloid cell subsets^81^ and we showed expression in BMPs (CMPs and GMPs).

Our findings suggest that gut microbiota translocation induces TI of BMPs that can underlie pathologies related to low-grade systemic inflammation. The identification of *E. faecalis* translocation to the BM as a main driver of this effect and its sensing by Mincle receptor in BMPs offers potential targets for intervention in pathophysiological settings where gut permeability is altered.

## Acknowledgments

We are grateful to members of the D.S. laboratory for discussions and critical reading of the manuscript, particularly to Gillian Dunphy. We thank the CNIC facilities and personnel for assistance. I.R-V is funded by (FJC2021-048099-I). P.B. was funded by grant BES-2014-069933 (“Ayudas para Contratos Predoctorales para la Formación de Doctores 2014”) from the Spanish Ministry of Economy, Industry and Competitiveness (MINECO). A.J-C is supported by an FPU Fellowship from the Spanish Ministry of Universities (FPU18/05434) and by a postgraduate fellowship of the City Council of Madrid at the Residencia de Estudiantes (2020–2021). M.M-L is supported by a Junior leader postdoctoral fellowship from “la Caixa” Foundation (116923). J.M. and C.G.C. are predoctoral fellows of MINECO and Junta de Andalucía, respectively. C.delF is funded by Instituto de Salud Carlos III through the project CP20/00106 and PI21-01178 (co-funded by European Union). JD laboratory is funded by Spanish Ministerio de Ciencia e Innovación PID2020-116347RB-I00/AEI/10.13039/501100011033. Work in the DS laboratory is funded by the CNIC; by the European Union’s Horizon 2020 research and innovation program under grant agreement ERC-2016-Consolidator Grant 725091; by Spanish Ministerio de Ciencia e Innovación PID2019-108157RB/AEI/; and CPP2021-008310/AEI/10.13039/501100011033 Unión Europea NextGenerationEU/PRTR; by Comunidad de Madrid (P2022/BMD-7333 INMUNOVAR-CM); by Atresmedia (Constantes y Vitales prize); by a research agreement with Inmunotek S.L.; by Fundació La Marató de TV3 (201723); and by “la Caixa” Foundation (LCF/PR/HR20/00075 and LCF/PR/HR22/00253). The CNIC is supported by the Instituto de Salud Carlos III (ISCIII), the MICINN and the Pro CNIC Foundation, and is a Severo Ochoa Center of Excellence (CEX2020-001041-S funded by MCIN/AEI /10.13039/501100011033). Work in the S.I. laboratory is funded by the Spanish Ministerio de Ciencia, Innovación (MICINN), Agencia Estatal de Investigación and Fondo Europeo de Desarrollo Regional, RTI2018-094484-B-I00; PID2021-125415OB-I00 and RYC-2016-19463.

## Author contributions

I.R-V., and D.S. conceived and designed the project and laboratory experiments; I.R-V, P.B, A.J-C, S.M-C, C.G-C, P.M-M, M.B-G, CM.D-R, J.M, M.M-L and S.I performed the laboratory experiments; IR-V., S.I., A.J-C and D.S. analyzed and interpreted the laboratory experiments; M.J.G., A.Q., A.D., E.P, L.C, M.M-L, C.delF and J.L.S performed essential analyses; J.D provided essential reagents; I.F-L provided essential support. I.R-V. and D.S wrote the manuscript. All authors revised and edited the manuscript prior to submission.

## Declaration of interests

J.L.S., L.C., CM.D-R and S.M-C. were employees of Inmunotek S.L. at the time of the work. The D.S. lab receives funds from a collaboration agreement between CNIC and Inmunotek. The other authors declare no competing interests.

## STAR Methods

### Experimental Animals

Mice were bred and maintained in groups of 5 animals per cage at the CNIC under specific pathogen-free conditions. Unless otherwise stated, males of 6-8 weeks old were used. Animal studies were approved by the local ethics committee. All animal procedures conformed to EU Directive 2010/63EU and Recommendation 2007/526/EC regarding the protection of animals used for experimental and other scientific purposes, enforced by Spanish law under Real Decreto 1201/2005. Colonies included C57BL/6J (Jackson Laboratory); C57BL/6J male germ-free (GF) mice (University of Granada, Granada, Spain); Clec4e^−/−^ (B6.Cg-Clec4etm1.1Cfg) mice were kindly provided by the Scripps Research Institute, through R. Ashman and C. Wells (Griffiths University, Australia)^82^; *Rag1*^−/−^ mice (B6.129S7-Rag1tm1Mom/J, The Jackson Laboratory).

### DSS models

6-8-week-old male mice were treated (DSS) or not (CTR) in drinking water with dextran sulfate sodium (DSS) 36–55 KDa 3% w/v for 5 days, after a 5-day rest the mice were sacrificed. All mice were fed with standard chow for the whole experiment. The disease activity index (DAI) was evaluated after 10 days for each animal according to the scores listed in the Table below. Where indicated, mice were co-treated in the drinking water with antibiotics-fungizone cocktail (Vancomycin (0.5g/L), Metronidazole (1g/L), Neomycin (1g/L), Ampicillin(1g/L) and Amphotericin B (1mg/L).

| SCORE | WEIGHT | Blood in feces | Stool formation |
| --- | --- | --- | --- |
| 0 | no change | - | normal |
| 1 | decrease by 1% | + | soft but formed |
| 2 | decrease by 3% | Blood visible | very soft |
| 3 | decrease by 5% | Bleeding out | diarrhea |

### Isolation and *ex vivo* stimulation of murine bone marrow cells

Bone marrow was isolated from femora and tibiae of 8- to -10-week-old male mice on WT C57BL/6J background kept in specific pathogen free (SPF) or Germ-free (GF) conditions, or *Clec4e*^−/−^, or *Rag1*^−/−^ kept in SPF. Bone marrow cell suspensions were prepared as described^83^. Briefly, bones were flushed with Roswell Park Memorial Institute medium (RPMI)-1640 supplemented with 10% Fetal Bovine Serum (FBS), 2mM L-glutamine, 100μg/ml penicillin, 100μg/ml streptomycin (all from Gibco), 10mM HEPES (Thermo Fisher), 1nM sodium pyruvate (Lonza), 100μM non-essential amino acids, herein referred to as “Complete RPMI”. Cells were centrifuged at 4 °C (5 min; 350xg). Erythrocytes were lysed by resuspending the pellet in red blood cell lysis buffer (150mM NH_4_Cl; 10mM KHCO_3_; 130μM ethylenediaminetetraacetic acid (EDTA)). The reaction was stopped by adding RPMI and cells were again centrifuged and used directly. For ex vivo stimulation, total cells from BM were cultured in a 96-well flat-bottom plate (10^5^ cells/well) in complete RPMI medium. After a 2-hour resting period in the incubator (37°C, 5% CO_2_), cells were stimulated with TLR-ligands for 6 hours: LPS (10ng/ml; TLR4 ligand), Pam3CSK4 (10ng/ml; TLR1/2 ligand), and CpG (10μM; TLR9 ligand) and supernatants were collected for cytokine analysis.

### Cell lines

The L929 cell line (ATCC® CCL-1TM), used to produce M-CSF supernatant, was grown on 175 cm^2^ cell culture flasks (Stemcell) and resuspended in Complete RPMI minus HEPES. Supernatants were obtained by filtering L929 culture supernatant at day 15 through 0.22 μm Stericup Filter units (Merck Millipore) and were used to subsequently supplement the medium for the generation of bone marrow derived macrophages.

Parental B3Z cells were kindly provided by N. Shastri, University of California) and express a β-gal reporter for nuclear factor of activated T cells (NFAT)^84^. Parental B3Z cells were transduced with retroviruses encoding a chimera of the extracellular domain of either mouse Mincle, Dectin-1 or Dectin-2 fused to the transmembrane region from NKRP1B and the intracellular tail of CD3ζ followed by an internal ribosome entry site (IRES) sequence and the GFP gene^85^. In addition, B3Z cells were transduced with retroviruses encoding for FcRγ chain, Syk and mouse Mincle, to test signaling through the natural receptor^42^. Binding of ligands can be detected by NFAT reporter activation and induction of β-gal activity^86^.

### Differentiation, culture, and stimulation of BMDM

Murine BMDMs were generated as previously described^87^ with some modifications. BM cells from WT C57BL/6J, *Clec4e* ^−/−^, *Rag1^−/−^*, or C57BL/6J Germ-free mice, were cultured in Complete RPMI supplemented with 30% M-CSF (obtained from L929 cell supernatants) during 5 days in sterile, but not tissue-culture treated, 10cm Petri dishes. BMDM, obtained from CTR or DSS pretreated mice, were plated in equal numbers (10^5^ cells per well) in 96-well plates (200μl final volume, Corning) and rested overnight. BMDMs were then stimulated with TLR-ligands for 24 hours: LPS (10ng/ml; TLR4 ligand), Pam3CSK4 (10ng/ml; TLR1/2 ligand) or CpG (10μM; TLR9 ligand). Subsequently, supernatants were collected for cytokine analysis. In separate experiments, BMDMs obtained from C57BL/6 mice were primed with heat killed (80°C-15min) HK-EF (2×10^2^ CFU/mL) for 24 hours. In the indicated experiments, BMDMs were pre-treated with 5′-methylthioadenosine (MTA) (500 μM), rapamycin (10nM), Syk Inhibitor (R406 (1μM)) or Card9 inhibitor (BRD5529 (10 μM)) one hour before HK-EF addition.

### Human samples

#### Peripheral blood mononuclear cells isolation

Healthy donors were recruited from the Blood Donor Service of La Paz University Hospital. Peripheral blood mononuclear cells (PBMCs) from healthy donors were isolated from venous blood collected in EDTA-containing tubes using Ficoll-Plus (GE Healthcare Bio-Sciences) solution according to the manufacturer’s instructions. PBMCs were washed twice with phosphate buffer saline (PBS), counted using Trypan blue staining, and resuspended at 2×10^6^ cell/mL in RPMI-1640 (Gibco) supplemented with 0.01% penicillin/streptomycin (p/s) (Thermofisher; Waltham, MA, United States) and with 10% FBS (Gibco). PBMCs (6×10^6^ cells) were plated in 6-well plates with RPMI supplemented with 10% FBS and 0.01% penicillin/streptomycin (p/s) (Thermofisher; Waltham, MA, United States) and rested an overnight before stimulation.

#### Trained immunity protocol *in vitro*

HK-EF (2×10^2^ CFU/mL) or particulate β-glucan (5μg/mL) were added to the cell culture medium for 24 hours to induce training, cells were then washed with PBS and fresh media was added. 5 days post-training, the medium was removed and LPS were added in Complete RPMI, and supernatant was collected 24 hours after.

#### Trained immunity protocol *in vivo*

HK-EF (2×10^2^ CFU) or PBS were injected i.v at day 0. After 10 days, 4 weeks or 8 weeks, mice were challenge i.p with LPS 5μg. In other set of experiments a dose-response curve was performed with a range of doses of HK-EF between 2×10^1^-2×10^7^ and, after 10 days, mice were challenged i.p with LPS 5μg. One hour after LPS challenge blood was obtained and TNFα was measured by ELISA using Mouse TNFα DuoSet; R&D Systems in plasma according with the manufacture instruction.

#### Flow cytometry analysis of Bone marrow progenitors

For the analysis of bone marrow progenitor cells, single cell suspensions from BM were collected as described above. Cells were labelled with a viability staining Aqua dead cell stain kit (Thermofisher) 30 min. Then a cocktail of biotinylated antibodies against lineage (lin) markers CD3, B220, CD45, Cd11b (1:100), Gr-1 and TER-119 (1:20) all from BD Pharmigen for 20 minutes. Then, cells were additionally stained with MHCII-FITC (BD Pharmigen) ; CD115-PerCP (Biolegend); CD45.2-V450 (BD Biosciences); c-kit-PE (Biolegend); CD11c-BV650 (Biolegend); CD150-BV605 (Biolegend); Strep-BV711 (Biolegend); Sca1-PECy7 (Biolegend); CD135-APC (Biolegend); CD48-APCCy7 (Biolegend) (1:200) for 20 min. CMPs were identified as as lin− Sca1− c-kit+ CD48+ CD150+ ; GMPs were identified as lin− Sca1− c-kit+ CD48+ CD150-; MPP3 cells were identified as lin-Sca1+ c-kit+ CD135-CD150-CD48+. Samples were analyzed on the LSR Fortessa (BD), and quantification was performed using FlowJo software (TreeStar Inc).

#### mTOR activation by Phospho-S6 staining

Cultured BMDM from C57BL/6 or *Clec4e*^−/−^ mice were plated in equal number (6×10^5^ cells per well) in 6-well plates (2000-μl final volume, Corning) and rested overnight. Cells were treated in vitro with HK-EF 2×10^2^ CFU/mL, TDB 10 μg/mL or PBS (Ctrol). Some cells were also co-incubated with Rapamycin (10nM). After 24h cells were collected and labelled with Phospho-S6-PE (Cell Signalling #5316) according to the manufacture protocol, activation was measured by flow cytometry MFI.

#### Quantification of cytokines

Cell culture supernatants and plasma samples were used to analyze cytokine production. TNFα measurement by ELISA using Mouse TNFα DuoSet; R&D Systems to BMDM or plasma from mice and Human TNF-alpha DuoSet ELISA; R&D Systems to PBMC following manufacturer’s instructions.

#### Bone marrow graft

WT C57BL/6J CD45.1^+^ mice were irradiated with two doses of 550 rads (5.5 Gy), separated by at least 3 hours, for a total radiation dose of 1100 rads. After 2 hours, irradiated mice received BM cells extracted from the medulla of CD45.2+ treatment group or control mice 1:1. Grafted mice received at least 2×10^6^ BM cells in 100 ul PBS or 5×10^5^ GMP cells by i.v. injection, and antibiotics (100 mg/mice Ampicillin) were injected subcutaneously. To isolate GMP cells, BM cells were stained as was describe above and GMP were identified as Lin-c-kit+sca-1-CD48+CD150-cells. After 30 days the reconstitution efficiency was checked in blood by analyzing CD45.1 vs CD45.2 expression. After 60 days the mice were challenged in vivo with LPS 5μg and sacrificed after 1 hour. Blood was collected in EDTA-containing tubes to obtain plasma for cytokine analysis. Some mice were not injected with LPS to collect bone marrow as described above to analyze bone marrow progenitor cells.

#### Analysis of bacterial translocation

Liver and BM from DSS-treated, ABX-treated, or untreated mice were homogenized in endotoxin-free PBS with 0.1% Triton X-100 to release intracellular bacteria. After a short centrifugation, 100μl supernatant from 1mL was added to a LB agar plate (Hardy Diagnostics) and cultured at 37°C for 24 hours in aerobic or anaerobic conditions. The number of colonies was counted and calculated as colony-forming units (CFU) per ml.

#### Germ-free mice inoculation

Male Germ-free mice were colonized with whole gut microbiota from conventional C57BL/6. For this, fresh feces were collected, resuspended in sterile PBS 1/20 w/v, centrifuged at 30xg for 10 minutes and keep -80°C until use)^88^ Mice colonized were maintained in separate isolators for the duration of the experiment.

#### Germ-free mice mono-colonization

Isolated *E. faecalis cultured* as described above, was used to mono-colonize male Germ-free mice. The colonization was performed by oral gavage (100μl) for two consecutive days with 10^8^ CFU of live bacteria. 24 hours after the second dose, DSS treatment was carried out as was described above. Mice colonized were maintained in separate isolators for the duration of the experiment.

#### Bacteria identification

Bacterial identification was performed using several tests. Isolated bacteria were grown in Columbia 5% SG horse (Biomerieux), PVX Chocolate polyvitex, (Biomerieux) Mac Conkey, (Biomerieux) GBS modified agar medium (VWR) and Aesculin agar (Oxoid). After 24 hours incubation at 37°C 5% CO_2_, grown bacteria were analyzed by microbiological test, Gram staining (allows bacteria to be differentiated according to their cell wall) and catalase test following standard procedures. After, API 20 strep test (Biomerieux) was carried out following manufacturer’s instructions.

The fermentation tests are inoculated with an enriched medium which rehydrates the sugar substrates. Fermentation of carbohydrates is detected by a shift in the pH indicator. The reactions are read according to the Reading Table (included in the protocol) and the identification is obtained by referring to the Analytical Profile Index or using the identification software.

To confirm the bacterial species, Matrix-Assisted Laser Desorption Ionization Time-of-Flight (MALDI-TOF) was used. MALDI-TOF is a mass spectrometry technique to assess the unique “molecular fingerprint” of an organism. Briefly, an isolated colony was transferred with an inoculation loop and placed in a well of the plate, once it was dry, 1μl of 70% formic acid was added. When the sample was dry, 1μl of Bruker matrix solution (α-Cyano-4-hydroxycinnamic acid, HCCA) was added on the sample, and once dry, the plate was inserted into the MALDI BioTyper (Bruker Daltonics) for the identification. Spectra was analyzed by the BioTyper program (Bruker Daltonics, Inc) utilizing both the Bruker database and a previously created Inmunotek in-house database. The MALDI-TOF Biotyper system (Bruker Daltonics) provides a numerical score for the interpretation of results. Scores are classified globally into several categories. According to the manufacturer, the score thresholds for microorganism identification are currently as follows: a score of ≥1.80 represents high confidence and it is accepted for bacterial identification. The bacterial test standard (BTS) will be included as a positive quality control for all experiments.

#### Enterococcus faecalis culture

Isolated *E. faecalis* as was described above from BM of DSS treated mice, was grown in aerobic condition overnight 37°C in LB medium under agitation 200 rpm. Grown bacteria were pelleted at 8000 g for 10 min and quantified by optical density and serial dilution 1:100 on Agar LB.

### Seahorse assay

Real-time Extracellular Acidification Rate (ECAR) in BMDMs were determined with an XF-96 Extracellular Flux Analyzer (Seahorse Bioscience). BMDMs were plated in Seahorse Cell Culture Microplates at a density of 100,000 cells per well and left to attach overnight. In the case of TDB stimulated BMDMs, 10mg TDB was coated to the Seahorse plate 24 hours before BMDMs were plated. After 24 hours of TDB or HK-EF stimulation, culture media was replaced by Seahorse Assay Medium (DMEM, 100μg/ml penicillin, 100μg/mL streptomycin, 2mM glutamine, 25mM glucose, 1mM pyruvate) at pH 7.4. Four technical replicates were used for each condition.

Seahorse cartridges were hydrated overnight in distilled water and calibrated for 1 hour in Seahorse XF Calibrant solution in a non-CO_2_-corrected incubator at 37°C. Seahorse assay was run using the Glycolysis Stress Test Protocol. For the Mito Stress Test the following compounds were added: Port A – oligomycin (2μM); Port B - carbonyl cyanide-4-(trifluoromethoxy) phenylhydrazone (FCCP) (0.5μM); Port C – rotenone + antimycin A (0.5μM). ECAR was analyzed at the beginning of the protocol with 3 measurements before inhibitor injection.

### Fungal infections

Fungal infection was performed in WT, *Rag1^−/−^* or *Clec4e*^−/−^ mice. 10 days before infection, mice were primed i.v. with HK*-*EF (2×10^3^ CFU) or PBS. After this time, mice were i.v. infected with 100 µl of *Candida albicans* (1.5×10^5^ CFU per mouse) or (7.5×10^4^ CFU per mouse in Rag1^−/−^), both diluted in sterile PBS, and. In a separate set of experiments mice were primed i.n. with HK-EF (2×10^6^ CFU). Mice were monitored for weight, general health, and survival, following institutional guidance. *Candida albicans* (strain SC5314, clinical isolate) was kindly provided by Prof. C. Gil (Complutense University, Madrid, Spain). The fungus was grown on yeast extract-peptone-dextrose (YPD)-agar plates (Sigma) at 30°C for 24 hours to maintain the degree of virulence. After 24 hours, the colony was grown in liquid YPD at 30°C for 24 hours under agitation. Fungal titer was quantified and resuspended in PBS for injection.

### Assay of B3Z cell lines

Parental or transduced B3Z cells were plated in 96-well plates and incubated with plated TDB/Zymosan/*Candida albicans* as positive control respectability. Cells were cultured in RPMI 1640 supplemented with 2 mM L-glutamine, 100 U/ml penicillin, 100 μg/ml streptomycin, 50 μM 2-mercaptoethanol, and 10% heat-inactivated fetal bovine serum (FBS) (all from Life Technologies, Carlsbad, CA) at 37°C. After overnight culture, cells were lysed in lysis buffer containing chlorophenol red-β-D-galactopyranoside (CPRG, Roche)-containing buffer. 1-4 hours later, LacZ activity was measured by absorbance, with O.D. 595 measured relative to a reference of O.D. 655 nm on a spectrophotometer (Benchmark Plus. Bio Rad).

### Human Fc recombinant assay

Mincle-hFc, generated as described^42^, and control-hFc (R&D Systems) were prepared in 10% skimmed milk in PBS and incubated overnight at 4°C on a rotating wheel, washed three times and labeled with PE-conjugated goat anti-hFc antibody (eBioscience). Specific binding of Mincle was detected by PE-positive signal by flow cytometry. Human Mincle-Fc chimera was blocked with anti-human Mincle 2F2 clone (Sigma-Aldrich). An isotype matched antibody (clone GC323, mouse IgM) was used as a negative control (Sigma Aldrich).

### ATAC-Sequencing

BM cells were isolated as described above. Lineage negative (Lin^−^) cells were enriched by staining with a panel of biotinylated antibodies against lineage committed cells (CD3, CD45R, CD11b (1:100) Ter-119, Gr-1 (1:20), all from BD Pharmigen). After washing, cells were incubated with streptavidin magnetic beads (Miltenyi) and passed through LS columns (Miltenyi), the negative fraction was collected for further staining with Sca-1 (APC), c-kit (CD117; PE), CD48 (APCCy7), CD150 (BV650), Streptavidin (BV421) (1:200) antibodies and with the viability marker Hoechst 33258 prior to sorting by FACS Aria II. GMPs were identified as Lin^−^c-kit^+^ sca-1^1−^ CD48^+^ CD150^−^ cells. 30,000-50,000 flow sorted GMPs were collected in ice-cold Flow cytometry buffer, and immediately treated as described previously^89^. In brief, cells were spun down in 25 μl cold lysis buffer at 500g for 20 minutes at 4°C. Afterwards, the transposition reaction was started by adding Nextera’s Tn5 Transposase in reaction buffer. The transposition reaction mix was incubated for 30 minutes at 37°C, DNA was purified using a Qiagen MinElute PCR purification kit. ATAC-seq libraries were purified using a PCR purification MinElute kit (Qiagen). Afterward, AMPure XP beads (Beckman) were used for size selection of the libraries, which were subsequently quantified using a Qubit fluorometer (ThermoFisher Scientific). Libraries size and quality were assessed using a 2100 Bioanalyzer instrument (Agilent). Libraries were sequenced in paired-end reads (2×50) using a NextSeq 2000 System. (Illumina) at the Genomics Unit of the CNIC.

### ATAC-Seq data analysis

Sequencing reads were pre-processed by means of a pipeline that used FastQC, to asses read quality, and Cutadapt^90^ to trim sequencing reads, eliminating Illumina and Nextera transposase adapter contaminations, and to discard reads that were shorter than 30 bp. Resulting reads were then mapped against reference genome GRCm38/MM10 with a pipeline that used bowtie2^91^ as aligner, Piccard to mark duplicate alignments, and samtools^92^ to eliminate duplicate, chimeric and sub-optimally multi-mapped alignments, keeping only properly paired and mapped reads. Alignments against the mitochondrial genome or chromosome Y were also removed. The final number of correctly aligned, filtered read pairs was between 12 and 47 million. TSS enrichment values, calculated with HOMER’s annotatePeaks function^93^ had values between 31 and 39. Once filtered alignments had been obtained, peaks (accessible DNA regions) were called with MACS3 ^94^, using parameters “--nomodel --shift -100 --extsize 200”, and “-q 0.05” as the false discovery rate cut-off. The numbers of peaks detected were between 97 and 131 thousand. Next, filtered alignments and peaks, in BAM and XLS formats, respectively, were processed with the R package DiffBind to define a consensus set of 154,415 peaks, representing the complete collection of accesible DNA regions detected in any condition. DiffBind was also used to calculate and normalize peak coverage across samples, and to identify differentially accessible regions (DARs) in pair-wise contrasts, using EdgeR as analysis method. The fraction of reads in peaks (FRiP score), as calculated by DiffBind, was between 0.58 and 0.73. The number of DARs (with p < 0.01) was between 1,474 and 56,012, depending on the contrast. DAR collections were annotated with HOMER^92^ to calculate the association of peaks to various genomic features, to identify the closest gene for each peak and to perform functional enrichment analyses, and to identify enriched motifs. In order to perform additional functional enrichment analyses with GSEA ^95^, peaks in the consensus peak set were associated to the corresponding closest genes with HOMER. Then, unique gene-peak pairs were defined for each contrast by keeping those pairs with the maximal absolute ATAC-Seq accesibility logFC value, as long as gene-peak distance was shorter than 1MB. Gene lists were finally used to perform functional enrichment analyses with GSEA preranked, using accesibility logFC values to sort gene lists. The association between DARs and promoters and proximal and distal enhancers as defined in ENCODE’s candidate Cis-Regulatory Element collection (cCRE; http://genome.ucsc.edu/cgi-bin/hgTrackUi?db=hg38&g=encodeCcreCombined) was tested with GAT ^96^, Genomic heatmaps were generated with deepTools2^97^. Other data manipulations and graphical representations were produced with R.

### DDA:TDB liposomes

Liposomes were prepared as described previously^98^. Briefly, a 5mg/mL solution of dimethyl-dioctadecyl-ammonium bromide (DDA) and TDB (Trehalose-6,6-dibehenate) were prepared using as dissolvent chloroform/methanol (9:1 v/v). DDA solution (5mg/mL) and TDB solution (5mg/mL) were mixed in a ratio 5:1 w/w (DDA:TDB). The organic solvent was removed using a roto-evaporator followed by flushing with N_2_ to form a thin lipid film on the bottom of a round-bottomed flask. The DDA:TDB complex was stored at -20°C. The lipid film was hydrated with 2ml of PBS to a final concentration of 2 mg/mL DDA and 0.25 mg/mL TDB added in the case of DDA:TDB, by heating for 20 min at 60 °C and used immediately to inject 10μg per mice.

### Statistical analysis

Mice were included in the studies in a blind manner and randomly assigned to receive or not DSS (1:1 simple randomization). The same procedure was used to allocate mice to different groups to be treated with HK-EF or PBS and DDA:TDB or DDA.

Sample size was calculated according to a previously performed pilot study to determine the DSS, and the infective dose of *C. albicans*. For the *C. albicans* experiments, we considered two independent study groups with a dichotomous primary endpoint represented by mortality. As statistical parameters we chose an anticipated incidence of 50% for one group and 20% for another one, an alpha error of 0.05 and a power of 0.8. In the experimental models, the distribution normality of data was evaluated with the Kolmogorov-Smirnov (KS) test. Differences in weight loss between HK-EF and PBS groups were compared using a two-way ANOVA test, followed by multiple comparisons corrected using Bonferroni statistical hypothesis testing. Survival curves were compared with log rank (Mantel-Cox) test. Two-tailed unpaired Student’s t test was generally used to evaluate statistical significance between two conditions, except for experiments *in vitro* with BMDM and human monocytes in which paired Student’s t test was applied. Differences were considered significant at P < 0.05 (∗p < 0.05; ∗∗p < 0.01; ∗∗∗p < 0.005; ****< 0.001). (Prism 9, Graph-Pad Software). The statistical test used and the definition of n are indicated in figure legends.

## Data and code availability

- ATAC-Seq data have been deposited at NCBI’s Gene Expression Omnibus and are publicly available as of the date of publication. Accession number is listed in the key resources table.
- This paper does not report original code.
- Any additional information required to reanalyze the data reported in this work paper is available from the Lead Contact upon request

---

**Fig. S1.**
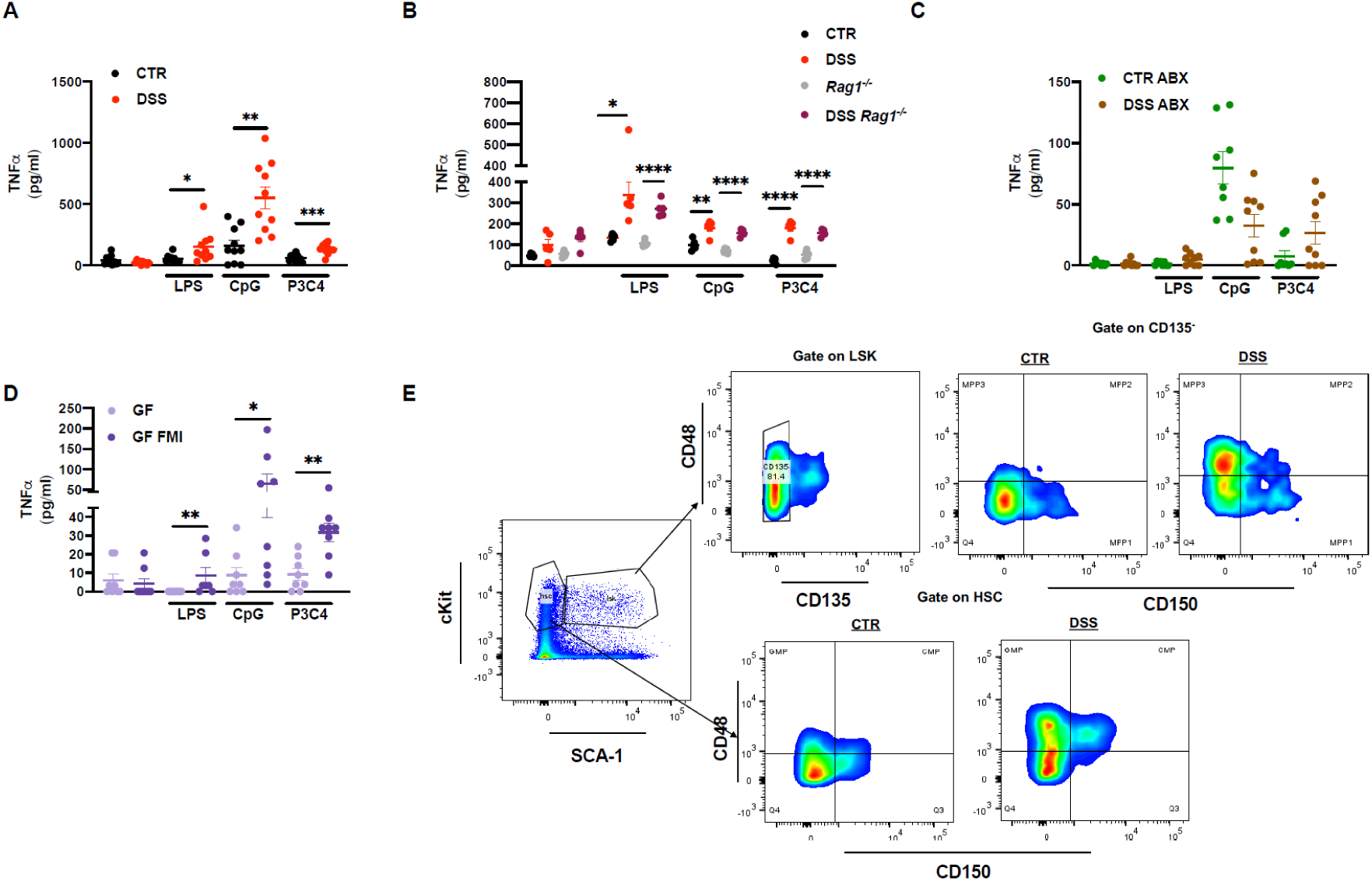
Gut microbiota translocation induces trained immunity in myeloid bone marrow progenitors. Related to Figure 1. (A) Mice were treated or not with DSS as indicated in the outline of Fig. 1A, total bone marrow (BM) was extracted, plated and stimulated 24h with LPS, CpG or P3C4. TNFα production measured by ELISA in culture supernatant of BM from indicated treatments and stimulations. Two pooled independent experiments (n=10). (B) WT and *Rag1*^−/−^ mice were treated or not with DSS as indicated in the outline of Fig. 1A, BM extracted, BMDMs generated and stimulated with LPS, CpG or P3C4 for 24h. TNFα production measured by ELISA in culture supernatant of BMDMs from indicated treatments and stimulations. One representative experiment, n=5. (C) Mice treated as described in A, but with or without antibiotics (ABX) and BM cells stimulated as in A, TNFα production measured by ELISA in culture supernatant of BM from indicated treatments and stimulations. Two pooled independent experiments (n=8-9). (D) Germ-free (GF) mice inoculated or not with fecal microbiota (FMI) were treated with DSS as indicated in the outline of Fig. 1D, total bone marrow (BM) was extracted, plated and stimulated 24h with LPS, CpG, or P3C4. TNFα production measured by ELISA in culture supernatant of BM from indicated treatments and stimulations. Two pooled independent experiments (n=8). (E) Gating strategy for the analysis of myeloid BMPs. Individual data and mean ± SEM is shown unless otherwise stated. Unpaired Student’s t test was used to compare groups; *p < 0.05; **p < 0.01; *** p < 0.005; ****p < 0.001.

**Fig. S2.**
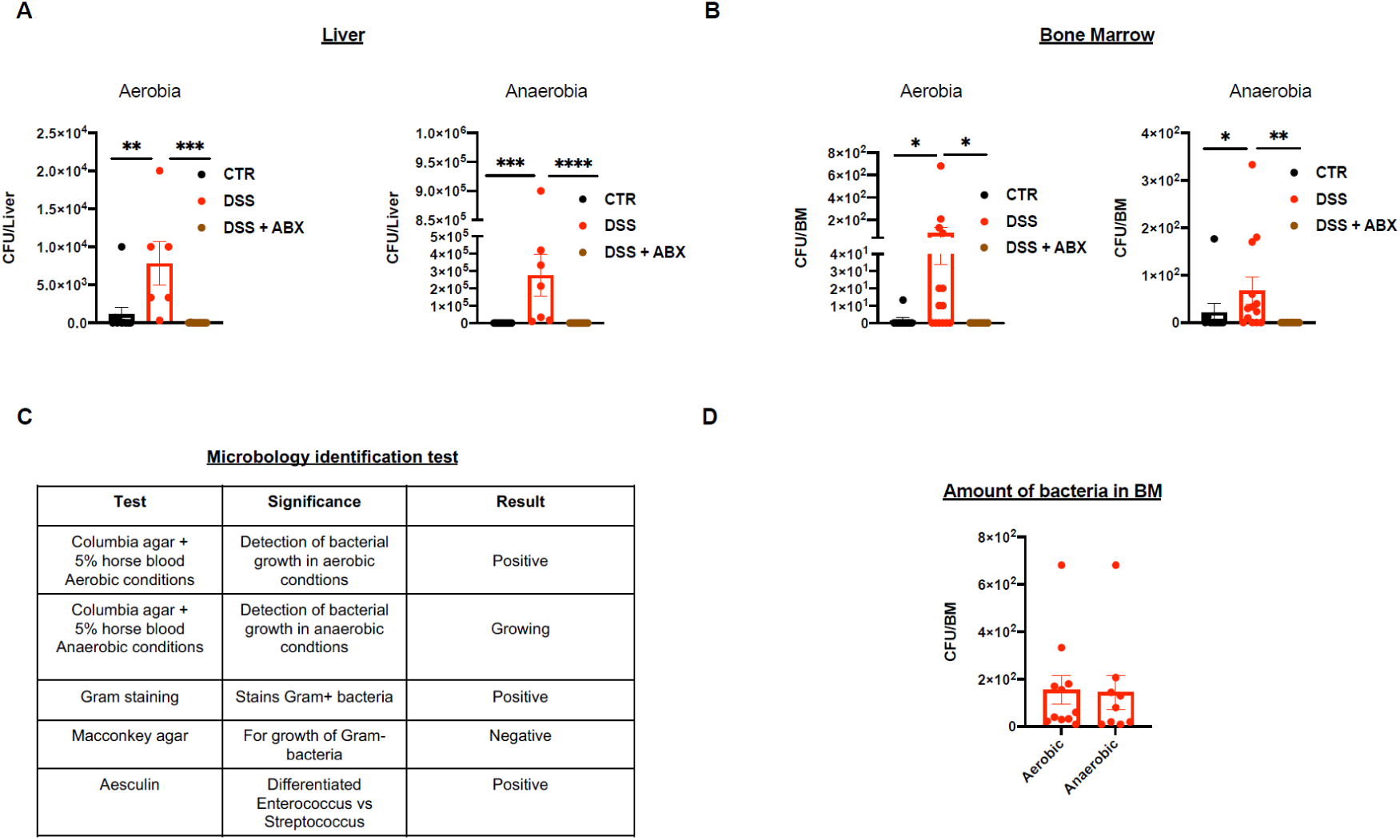
Analysis of bacterial translocation. Related to Figure 2. (A, B) Bacterial translocation in liver, n=7-10 (A) and BM, n= 7-14 (B) as determined by quantification of colony forming units in aerobic (left) and anaerobic (right) non-selective cultures. Three pooled independent experiments. (C) Results of microbiology test performed in cultured bacteria isolated from BM of DSS treated mice. Three independent experiments. (D) Quantification of bacteria in BM of DSS-treated mice, determined by colony growth from BM flushed in PBS 0.1% Triton in aerobic and anaerobic non-selective cultures, considering only the mice with positive translocation. Individual data and mean ± SEM is shown unless otherwise stated. Unpaired Student’s t test was used to compare groups; *p < 0.05; **p < 0.01; *** p < 0.005; ****p < 0.001.

**Fig S3.**
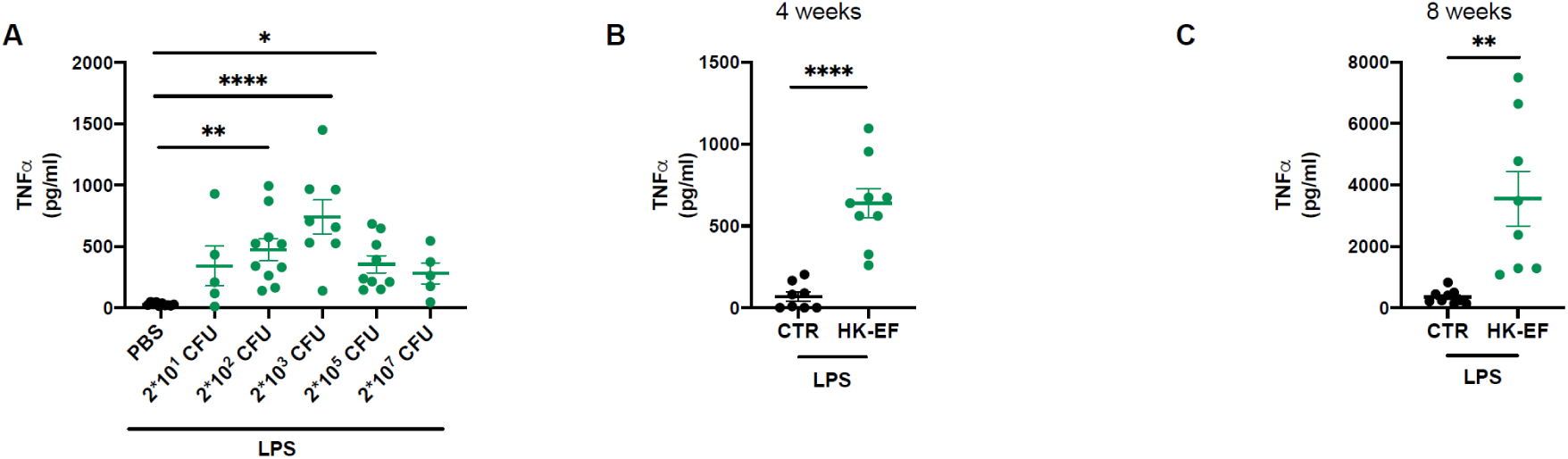
Dose response and timing. Related to Figure 3. A) TNFα concentration in plasma from mice treated with increasing doses of HK-EF in vivo (2×10^1^-2×10^7^ CFU), and after 10 days, challenged for 1h with intraperitoneal LPS. Two pooled independent experiments, n=5-10 (B, C) TNFα concentration in plasma from mice primed with *E. faecalis* in vivo with 2×10^2^ CFU and challenged with LPS i.p. after 4 weeks (B) or 8 weeks (C) later. Two pooled independent experiments, n=8-9 Individual data and mean ± SEM is shown unless otherwise stated. Unpaired Student’s t test was used to compare groups; *p < 0.05; **p < 0.01; *** p < 0.005; ****p < 0.001.

**Fig S4.**
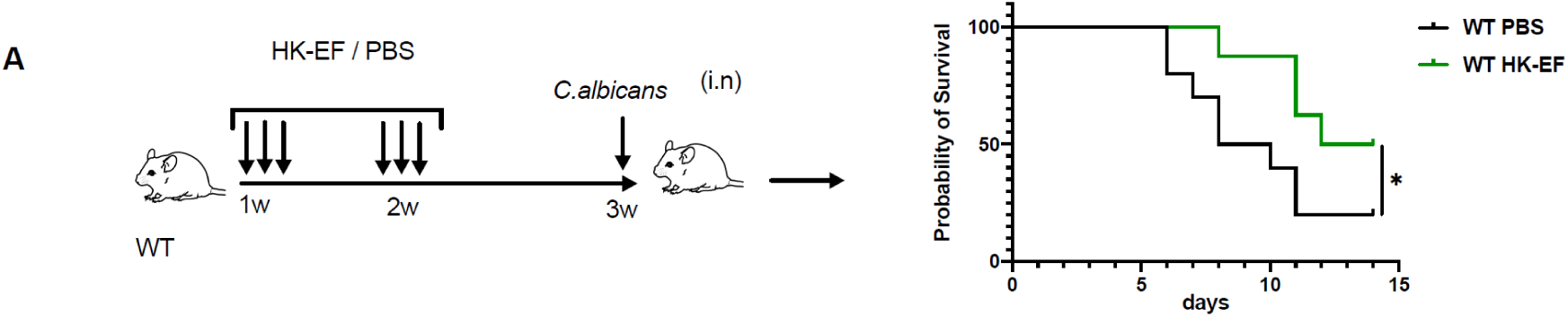
HK-EF Intranasal protection. Related to Figure 6. (A) WT mice were treated or not in with HK-EF for 2 weeks and infected with *C. albicans* the following week as indicated in the outline (left) The survival curve is shown (right). Results from a pool of two independent experiments are shown, n=10 for PBS, n=8 for HK-EF. Survival curves were compared with log rank (Mantel-Cox); *p < 0.05.

## References

1. Mowat, A.M. (2018). To respond or not to respond - a personal perspective of intestinal tolerance. Nat. Rev. Immunol. 18, 405–415. 10.1038/s41577-018-0002-x.

2. Mousa, W.K., Chehadeh, F., and Husband, S. (2022). Microbial dysbiosis in the gut drives systemic autoimmune diseases. Front. Immunol. 13, 906258. 10.3389/fimmu.2022.906258.

3. Sonnenberg, G.F., Monticelli, L.A., Alenghat, T., Fung, T.C., Hutnick, N.A., Kunisawa, J., Shibata, N., Grunberg, S., Sinha, R., Zahm, A.M., et al. (2012). Innate lymphoid cells promote anatomical containment of lymphoid-resident commensal bacteria. Science 336, 1321–1325. 10.1126/science.1222551.

4. Robles-Vera, I., Toral, M., de la Visitación, N., Sánchez, M., Gómez-Guzmán, M., Muñoz, R., Algieri, F., Vezza, T., Jiménez, R., Gálvez, J., et al. (2020). Changes to the gut microbiota induced by losartan contributes to its antihypertensive effects. Br. J. Pharmacol. 177, 2006–2023. 10.1111/bph.14965.

5. Okumura, R., and Takeda, K. (2018). Maintenance of intestinal homeostasis by mucosal barriers. Inflamm. Regen. 38, 5. 10.1186/s41232-018-0063-z.

6. Alexander, J.W., Boyce, S.T., Babcock, G.F., Gianotti, L., Peck, M.D., Dunn, D.L., Pyles, T., Childress, C.P., and Ash, S.K. (1990). The process of microbial translocation. Ann. Surg. 212, 492–496. 10.1097/00000658-199010000-00012.

7. Berthelot, J.-M., and Wendling, D. (2020). Translocation of dead or alive bacteria from mucosa to joints and epiphyseal bone-marrow: facts and hypotheses. Jt. bone spine 87, 31–36. 10.1016/j.jbspin.2019.01.004.

8. Manfredo Vieira, S., Hiltensperger, M., Kumar, V., Zegarra-Ruiz, D., Dehner, C., Khan, N., Costa, F.R.C., Tiniakou, E., Greiling, T., Ruff, W., et al. (2018). Translocation of a gut pathobiont drives autoimmunity in mice and humans. Science 359, 1156–1161. 10.1126/science.aar7201.

9. Macpherson, A.J., and Uhr, T. (2004). Induction of protective IgA by intestinal dendritic cells carrying commensal bacteria. Science 303, 1662–1665. 10.1126/science.1091334.

10. Hand, T.W., Dos Santos, L.M., Bouladoux, N., Molloy, M.J., Pagán, A.J., Pepper, M., Maynard, C.L., Elson, C.O. 3rd, and Belkaid, Y. (2012). Acute gastrointestinal infection induces long-lived microbiota-specific T cell responses. Science 337, 1553–1556. 10.1126/science.1220961.

11. Sandler, N.G., and Douek, D.C. (2012). Microbial translocation in HIV infection: causes, consequences and treatment opportunities. Nat. Rev. Microbiol. 10, 655–666. 10.1038/nrmicro2848.

12. Costa, F.R.C., Françozo, M.C.S., de Oliveira, G.G., Ignacio, A., Castoldi, A., Zamboni, D.S., Ramos, S.G., Câmara, N.O., de Zoete, M.R., Palm, N.W., et al. (2016). Gut microbiota translocation to the pancreatic lymph nodes triggers NOD2 activation and contributes to T1D onset. J. Exp. Med. 213, 1223–1239. 10.1084/jem.20150744.

13. Daillère, R., Vétizou, M., Waldschmitt, N., Yamazaki, T., Isnard, C., Poirier-Colame, V., Duong, C.P.M., Flament, C., Lepage, P., Roberti, M.P., et al. (2016). Enterococcus hirae and Barnesiella intestinihominis Facilitate Cyclophosphamide-Induced Therapeutic Immunomodulatory Effects. Immunity 45, 931–943. 10.1016/j.immuni.2016.09.009.

14. Meisel, M., Hinterleitner, R., Pacis, A., Chen, L., Earley, Z.M., Mayassi, T., Pierre, J.F., Ernest, J.D., Galipeau, H.J., Thuille, N., et al. (2018). Microbial signals drive pre-leukaemic myeloproliferation in a Tet2-deficient host. Nature 557, 580–584. 10.1038/s41586-018-0125-z.

15. Viaud, S., Saccheri, F., Mignot, G., Yamazaki, T., Daillère, R., Hannani, D., Enot, D.P., Pfirschke, C., Engblom, C., Pittet, M.J., et al. (2013). The intestinal microbiota modulates the anticancer immune effects of cyclophosphamide. Science 342, 971–976. 10.1126/science.1240537.

16. Danne, C., Ryzhakov, G., Martínez-López, M., Ilott, N.E., Franchini, F., Cuskin, F., Lowe, E.C., Bullers, S.J., Arthur, J.S.C., and Powrie, F. (2017). A Large Polysaccharide Produced by Helicobacter hepaticus Induces an Anti-inflammatory Gene Signature in Macrophages. Cell Host Microbe 22, 733–745.e5. 10.1016/j.chom.2017.11.002.

17. Gasaly, N., de Vos, P., and Hermoso, M.A. (2021). Impact of Bacterial Metabolites on Gut Barrier Function and Host Immunity: A Focus on Bacterial Metabolism and Its Relevance for Intestinal Inflammation. Front. Immunol. 12, 658354. 10.3389/fimmu.2021.658354.

18. Zheng, D., Liwinski, T., and Elinav, E. (2020). Interaction between microbiota and immunity in health and disease. Cell Res. 30, 492–506. 10.1038/s41422-020-0332-7.

19. Negi, S., Das, D.K., Pahari, S., Nadeem, S., and Agrewala, J.N. (2019). Potential Role of Gut Microbiota in Induction and Regulation of Innate Immune Memory. Front. Immunol. 10, 2441. 10.3389/fimmu.2019.02441.

20. Netea, M.G., Quintin, J., and van der Meer, J.W.M. (2011). Trained immunity: a memory for innate host defense. Cell Host Microbe 9, 355–361. 10.1016/j.chom.2011.04.006.

21. Netea, M.G., Joosten, L.A.B., Latz, E., Mills, K.H.G., Natoli, G., Stunnenberg, H.G., O’Neill, L.A.J., and Xavier, R.J. (2016). Trained immunity: A program of innate immune memory in health and disease. Science 352, aaf1098. 10.1126/science.aaf1098.

22. Krautkramer, K.A., Kreznar, J.H., Romano, K.A., Vivas, E.I., Barrett-Wilt, G.A., Rabaglia, M.E., Keller, M.P., Attie, A.D., Rey, F.E., and Denu, J.M. (2016). Diet-Microbiota Interactions Mediate Global Epigenetic Programming in Multiple Host Tissues. Mol. Cell 64, 982–992. 10.1016/j.molcel.2016.10.025.

23. Miro-Blanch, J., and Yanes, O. (2019). Epigenetic Regulation at the Interplay Between Gut Microbiota and Host Metabolism. Front. Genet. 10, 638. 10.3389/fgene.2019.00638.

24. Schulthess, J., Pandey, S., Capitani, M., Rue-Albrecht, K.C., Arnold, I., Franchini, F., Chomka, A., Ilott, N.E., Johnston, D.G.W., Pires, E., et al. (2019). The Short Chain Fatty Acid Butyrate Imprints an Antimicrobial Program in Macrophages. Immunity 50, 432–445.e7. 10.1016/j.immuni.2018.12.018.

25. Burgess, S.L., Leslie, J.L., Uddin, J., Oakland, D.N., Gilchrist, C., Moreau, G.B., Watanabe, K., Saleh, M., Simpson, M., Thompson, B.A., et al. (2020). Gut microbiome communication with bone marrow regulates susceptibility to amebiasis. J. Clin. Invest. 130, 4019–4024. 10.1172/JCI133605.

26. Luo, Y., Chen, G.-L., Hannemann, N., Ipseiz, N., Krönke, G., Bäuerle, T., Munos, L., Wirtz, S., Schett, G., and Bozec, A. (2015). Microbiota from Obese Mice Regulate Hematopoietic Stem Cell Differentiation by Altering the Bone Niche. Cell Metab. 22, 886–894. 10.1016/j.cmet.2015.08.020.

27. Clarke, T.B., Davis, K.M., Lysenko, E.S., Zhou, A.Y., Yu, Y., and Weiser, J.N. (2010). Recognition of peptidoglycan from the microbiota by Nod1 enhances systemic innate immunity. Nat. Med. 16, 228–231. 10.1038/nm.2087.

28. Deshmukh, H.S., Liu, Y., Menkiti, O.R., Mei, J., Dai, N., O’Leary, C.E., Oliver, P.M., Kolls, J.K., Weiser, J.N., and Worthen, G.S. (2014). The microbiota regulates neutrophil homeostasis and host resistance to Escherichia coli K1 sepsis in neonatal mice. Nat. Med. 20, 524–530. 10.1038/nm.3542.

29. Trompette, A., Gollwitzer, E.S., Yadava, K., Sichelstiel, A.K., Sprenger, N., Ngom-Bru, C., Blanchard, C., Junt, T., Nicod, L.P., Harris, N.L., et al. (2014). Gut microbiota metabolism of dietary fiber influences allergic airway disease and hematopoiesis. Nat. Med. 20, 159–166. 10.1038/nm.3444.

30. Mitroulis, I., Kalafati, L., Hajishengallis, G., and Chavakis, T. (2018). Myelopoiesis in the Context of Innate Immunity. J. Innate Immun. 10, 365–372. 10.1159/000489406.

31. van der Meer, J.W.M., Joosten, L.A.B., Riksen, N., and Netea, M.G. (2015). Trained immunity: A smart way to enhance innate immune defence. Mol. Immunol. 68, 40–44. 10.1016/j.molimm.2015.06.019.

32. Sancho, D., and Reis e Sousa, C. (2012). Signaling by myeloid C-type lectin receptors in immunity and homeostasis. Annu. Rev. Immunol. 30, 491–529. 10.1146/annurev-immunol-031210-101352.

33. Braganza, C.D., Teunissen, T., Timmer, M.S.M., and Stocker, B.L. (2017). Identification and Biological Activity of Synthetic Macrophage Inducible C-Type Lectin Ligands. Front. Immunol. 8, 1940. 10.3389/fimmu.2017.01940.

34. Martínez-López, M., Iborra, S., Conde-Garrosa, R., Mastrangelo, A., Danne, C., Mann, E.R., Reid, D.M., Gaboriau-Routhiau, V., Chaparro, M., Lorenzo, M.P., et al. (2019). Microbiota Sensing by Mincle-Syk Axis in Dendritic Cells Regulates Interleukin-17 and -22 Production and Promotes Intestinal Barrier Integrity. Immunity 50, 446–461.e9. 10.1016/j.immuni.2018.12.020.

35. Jensen, S.K., Pærregaard, S.I., Brandum, E.P., Jørgensen, A.S., Hjortø, G.M., and Jensen, B.A.H. (2022). Rewiring host-microbe interactions and barrier function during gastrointestinal inflammation. Gastroenterol. Rep. 10, goac008. 10.1093/gastro/goac008.

36. de la Visitación, N., Robles-Vera, I., Moleón, J., González-Correa, C., Aguilera-Sánchez, N., Toral, M., Gómez-Guzmán, M., Sánchez, M., Jiménez, R., Martin-Morales, N., et al. (2021). Gut Microbiota Has a Crucial Role in the Development of Hypertension and Vascular Dysfunction in Toll-like Receptor 7-Driven Lupus Autoimmunity. Antioxidants (Basel, Switzerland) 10. 10.3390/antiox10091426.

37. Saz-Leal, P., Del Fresno, C., Brandi, P., Martínez-Cano, S., Dungan, O.M., Chisholm, J.D., Kerr, W.G., and Sancho, D. (2018). Targeting SHIP-1 in Myeloid Cells Enhances Trained Immunity and Boosts Response to Infection. Cell Rep. 25, 1118–1126. 10.1016/j.celrep.2018.09.092.

38. Cheng, S.-C., Quintin, J., Cramer, R.A., Shepardson, K.M., Saeed, S., Kumar, V., Giamarellos-Bourboulis, E.J., Martens, J.H.A., Rao, N.A., Aghajanirefah, A., et al. (2014). mTOR- and HIF-1α-mediated aerobic glycolysis as metabolic basis for trained immunity. Science 345, 1250684. 10.1126/science.1250684.

39. Brandi, P., Conejero, L., Cueto, F.J., Martínez-Cano, S., Dunphy, G., Gómez, M.J., Relaño, C., Saz-Leal, P., Enamorado, M., Quintas, A., et al. (2022). Trained immunity induction by the inactivated mucosal vaccine MV130 protects against experimental viral respiratory infections. Cell Rep. 38, 110184. 10.1016/j.celrep.2021.110184.

40. Mata-Martínez, P., Bergón-Gutiérrez, M., and Del Fresno, C. (2022). Dectin-1 Signaling Update: New Perspectives for Trained Immunity. Front. Immunol. 13, 812148. 10.3389/fimmu.2022.812148.

41. Sancho, D., Joffre, O.P., Keller, A.M., Rogers, N.C., Martínez, D., Hernanz-Falcón, P., Rosewell, I., and Reis e Sousa, C. (2009). Identification of a dendritic cell receptor that couples sensing of necrosis to immunity. Nature 458, 899–903. 10.1038/nature07750.

42. Iborra, S., Martínez-López, M., Cueto, F.J., Conde-Garrosa, R., Del Fresno, C., Izquierdo, H.M., Abram, C.L., Mori, D., Campos-Martín, Y., Reguera, R.M., et al. (2016). Leishmania Uses Mincle to Target an Inhibitory ITAM Signaling Pathway in Dendritic Cells that Dampens Adaptive Immunity to Infection. Immunity 45, 788–801. 10.1016/j.immuni.2016.09.012.

43. Schoenen, H., Bodendorfer, B., Hitchens, K., Manzanero, S., Werninghaus, K., Nimmerjahn, F., Agger, E.M., Stenger, S., Andersen, P., Ruland, J., et al. (2010). Cutting edge: Mincle is essential for recognition and adjuvanticity of the mycobacterial cord factor and its synthetic analog trehalose-dibehenate. J. Immunol. 184, 2756–2760. 10.4049/jimmunol.0904013.

44. Martin-Cruz, L., Sevilla-Ortega, C., Benito-Villalvilla, C., Diez-Rivero, C.M., Sanchez-Ramón, S., Subiza, J.L., and Palomares, O. (2020). A Combination of Polybacterial MV140 and Candida albicans V132 as a Potential Novel Trained Immunity-Based Vaccine for Genitourinary Tract Infections. Front. Immunol. 11, 612269. 10.3389/fimmu.2020.612269.

45. Del Fresno, C., García-Arriaza, J., Martínez-Cano, S., Heras-Murillo, I., Jarit-Cabanillas, A., Amores-Iniesta, J., Brandi, P., Dunphy, G., Suay-Corredera, C., Pricolo, M.R., et al. (2021). The Bacterial Mucosal Immunotherapy MV130 Protects Against SARS-CoV-2 Infection and Improves COVID-19 Vaccines Immunogenicity. Front. Immunol. 12, 748103. 10.3389/fimmu.2021.748103.

46. Wirtz, S., Neufert, C., Weigmann, B., and Neurath, M.F. (2007). Chemically induced mouse models of intestinal inflammation. Nat. Protoc. 2, 541–546. 10.1038/nprot.2007.41.

47. Zeng, M.Y., Cisalpino, D., Varadarajan, S., Hellman, J., Warren, H.S., Cascalho, M., Inohara, N., and Núñez, G. (2016). Gut Microbiota-Induced Immunoglobulin G Controls Systemic Infection by Symbiotic Bacteria and Pathogens. Immunity 44, 647–658. 10.1016/j.immuni.2016.02.006.

48. Cani, P.D., Neyrinck, A.M., Fava, F., Knauf, C., Burcelin, R.G., Tuohy, K.M., Gibson, G.R., and Delzenne, N.M. (2007). Selective increases of bifidobacteria in gut microflora improve high-fat-diet-induced diabetes in mice through a mechanism associated with endotoxaemia. Diabetologia 50, 2374–2383. 10.1007/s00125-007-0791-0.

49. Chassaing, B., Koren, O., Goodrich, J.K., Poole, A.C., Srinivasan, S., Ley, R.E., and Gewirtz, A.T. (2015). Dietary emulsifiers impact the mouse gut microbiota promoting colitis and metabolic syndrome. Nature 519, 92–96. 10.1038/nature14232.

50. Christ, A., Günther, P., Lauterbach, M.A.R., Duewell, P., Biswas, D., Pelka, K., Scholz, C.J., Oosting, M., Haendler, K., Baßler, K., et al. (2018). Western Diet Triggers NLRP3-Dependent Innate Immune Reprogramming. Cell 172, 162–175.e14. 10.1016/j.cell.2017.12.013.

51. Wirtz, S., Popp, V., Kindermann, M., Gerlach, K., Weigmann, B., Fichtner-Feigl, S., and Neurath, M.F. (2017). Chemically induced mouse models of acute and chronic intestinal inflammation. Nat. Protoc. 12, 1295–1309. 10.1038/nprot.2017.044.

52. Mihaescu, A., Santen, S., Jeppsson, B., and Thorlacius, H. (2010). p38 Mitogen-activated protein kinase signalling regulates vascular inflammation and epithelial barrier dysfunction in an experimental model of radiation-induced colitis. Br. J. Surg. 97, 226–234. 10.1002/bjs.6811.

53. Edgar, L., Akbar, N., Braithwaite, A.T., Krausgruber, T., Gallart-Ayala, H., Bailey, J., Corbin, A.L., Khoyratty, T.E., Chai, J.T., Alkhalil, M., et al. (2021). Hyperglycemia Induces Trained Immunity in Macrophages and Their Precursors and Promotes Atherosclerosis. Circulation 144, 961–982. 10.1161/CIRCULATIONAHA.120.046464.

54. Mitroulis, I., Kalafati, L., Hajishengallis, G., and Chavakis, T. (2018). Myelopoiesis in the Context of Innate Immunity. J. Innate Immun. 10, 365–372. 10.1159/000489406.

55. Tortora, S.C., Bodiwala, V.M., Quinn, A., Martello, L.A., and Vignesh, S. (2022). Microbiome and colorectal carcinogenesis: Linked mechanisms and racial differences. World J. Gastrointest. Oncol. 14, 375–395. 10.4251/wjgo.v14.i2.375.

56. Roche-Lima, A., Carrasquillo-Carrión, K., Gómez-Moreno, R., Cruz, J.M., Velázquez-Morales, D.M., Rogozin, I.B., and Baerga-Ortiz, A. (2018). The Presence of Genotoxic and/or Pro-inflammatory Bacterial Genes in Gut Metagenomic Databases and Their Possible Link With Inflammatory Bowel Diseases. Front. Genet. 9, 116. 10.3389/fgene.2018.00116.

57. Llorente, C., Jepsen, P., Inamine, T., Wang, L., Bluemel, S., Wang, H.J., Loomba, R., Bajaj, J.S., Schubert, M.L., Sikaroodi, M., et al. (2017). Gastric acid suppression promotes alcoholic liver disease by inducing overgrowth of intestinal Enterococcus. Nat. Commun. 8, 837. 10.1038/s41467-017-00796-x.

58. Stothers, C.L., Burelbach, K.R., Owen, A.M., Patil, N.K., McBride, M.A., Bohannon, J.K., Luan, L., Hernandez, A., Patil, T.K., Williams, D.L., et al. (2021). β-Glucan Induces Distinct and Protective Innate Immune Memory in Differentiated Macrophages. J. Immunol. 207, 2785–2798. 10.4049/jimmunol.2100107.

59. Funes, S.C., Rios, M., Fernández-Fierro, A., Di Genaro, M.S., and Kalergis, A.M. (2022). Trained Immunity Contribution to Autoimmune and Inflammatory Disorders. Front. Immunol. 13, 868343. 10.3389/fimmu.2022.868343.

60. Hermans, L., De Pelsmaeker, S., Denaeghel, S., Cox, E., Favoreel, H.W., and Devriendt, B. (2021). β-Glucan-Induced IL-10 Secretion by Monocytes Triggers Porcine NK Cell Cytotoxicity. Front. Immunol. 12, 634402. 10.3389/fimmu.2021.634402.

61. Mitroulis, I., Ruppova, K., Wang, B., Chen, L.-S., Grzybek, M., Grinenko, T., Eugster, A., Troullinaki, M., Palladini, A., Kourtzelis, I., et al. (2018). Modulation of Myelopoiesis Progenitors Is an Integral Component of Trained Immunity. Cell 172, 147–161.e12. 10.1016/j.cell.2017.11.034.

62. Ishikawa, E., Ishikawa, T., Morita, Y.S., Toyonaga, K., Yamada, H., Takeuchi, O., Kinoshita, T., Akira, S., Yoshikai, Y., and Yamasaki, S. (2009). Direct recognition of the mycobacterial glycolipid, trehalose dimycolate, by C-type lectin Mincle. J. Exp. Med. 206, 2879–2888. 10.1084/jem.20091750.

63. Ishikawa, T., Itoh, F., Yoshida, S., Saijo, S., Matsuzawa, T., Gonoi, T., Saito, T., Okawa, Y., Shibata, N., Miyamoto, T., et al. (2013). Identification of distinct ligands for the C-type lectin receptors Mincle and Dectin-2 in the pathogenic fungus Malassezia. Cell Host Microbe 13, 477–488. 10.1016/j.chom.2013.03.008.

64. Del Fresno, C., Iborra, S., Saz-Leal, P., Martínez-López, M., and Sancho, D. (2018). Flexible Signaling of Myeloid C-Type Lectin Receptors in Immunity and Inflammation. Front. Immunol. 9, 804. 10.3389/fimmu.2018.00804.

65. Gong, W., Zheng, T., Guo, K., Fang, M., Xie, H., Li, W., Tang, Q., Hong, Z., Ren, H., Gu, G., et al. (2020). Mincle/Syk Signalling Promotes Intestinal Mucosal Inflammation Through Induction of Macrophage Pyroptosis in Crohn’s Disease. J. Crohns. Colitis 14, 1734–1747. 10.1093/ecco-jcc/jjaa088.

66. Sokol, H., Leducq, V., Aschard, H., Pham, H.-P., Jegou, S., Landman, C., Cohen, D., Liguori, G., Bourrier, A., Nion-Larmurier, I., et al. (2017). Fungal microbiota dysbiosis in IBD. Gut 66, 1039–1048. 10.1136/gutjnl-2015-310746.

67. Ni, J., Wu, G.D., Albenberg, L., and Tomov, V.T. (2017). Gut microbiota and IBD: causation or correlation? Nat. Rev. Gastroenterol. Hepatol. 14, 573–584. 10.1038/nrgastro.2017.88.

68. Kaser, A., Zeissig, S., and Blumberg, R.S. (2010). Inflammatory bowel disease. Annu. Rev. Immunol. 28, 573–621. 10.1146/annurev-immunol-030409-101225.

69. Li, M., Zhang, R., Li, J., and Li, J. (2022). The Role of C-Type Lectin Receptor Signaling in the Intestinal Microbiota-Inflammation-Cancer Axis. Front. Immunol. 13, 894445. 10.3389/fimmu.2022.894445.

70. Arts, R.J.W., Moorlag, S.J.C.F.M., Novakovic, B., Li, Y., Wang, S.-Y., Oosting, M., Kumar, V., Xavier, R.J., Wijmenga, C., Joosten, L.A.B., et al. (2018). BCG Vaccination Protects against Experimental Viral Infection in Humans through the Induction of Cytokines Associated with Trained Immunity. Cell Host Microbe 23, 89–100.e5. 10.1016/j.chom.2017.12.010.

71. Kaufmann, E., Sanz, J., Dunn, J.L., Khan, N., Mendonça, L.E., Pacis, A., Tzelepis, F., Pernet, E., Dumaine, A., Grenier, J.-C., et al. (2018). BCG Educates Hematopoietic Stem Cells to Generate Protective Innate Immunity against Tuberculosis. Cell 172, 176–190.e19. 10.1016/j.cell.2017.12.031.

72. Ziogas, A., and Netea, M.G. (2022). Trained immunity-related vaccines: innate immune memory and heterologous protection against infections. Trends Mol. Med. 28, 497–512. 10.1016/j.molmed.2022.03.009.

73. Vázquez, A., Fernández-Sevilla, L.M., Jiménez, E., Pérez-Cabrera, D., Yañez, R., Subiza, J.L., Varas, A., Valencia, J., and Vicente, A. (2020). Involvement of Mesenchymal Stem Cells in Oral Mucosal Bacterial Immunotherapy. Front. Immunol. 11, 567391. 10.3389/fimmu.2020.567391.

74. Sánchez-Ramón, S., Conejero, L., Netea, M.G., Sancho, D., Palomares, Ó., and Subiza, J.L. (2018). Trained Immunity-Based Vaccines: A New Paradigm for the Development of Broad-Spectrum Anti-infectious Formulations. Front. Immunol. 9, 2936. 10.3389/fimmu.2018.02936.

75. Martín-Cruz, L., Angelina, A., Baydemir, I., Bulut, Ö., Subiza, J.L., Netea, M.G., Domínguez-Andrés, J., and Palomares, O. (2022). Candida albicans V132 induces trained immunity and enhances the responses triggered by the polybacterial vaccine MV140 for genitourinary tract infections. Front. Immunol. 13, 1066383. 10.3389/fimmu.2022.1066383.

76. Jeyanathan, M., Vaseghi-Shanjani, M., Afkhami, S., Grondin, J.A., Kang, A., D’Agostino, M.R., Yao, Y., Jain, S., Zganiacz, A., Kroezen, Z., et al. (2022). Parenteral BCG vaccine induces lung-resident memory macrophages and trained immunity via the gut-lung axis. Nat. Immunol. 23, 1687–1702. 10.1038/s41590-022-01354-4.

77. Lorenzo-Gómez, M.F., Padilla-Fernández, B., García-Criado, F.J., Mirón-Canelo, J.A., Gil-Vicente, A., Nieto-Huertos, A., and Silva-Abuin, J.M. (2013). Evaluation of a therapeutic vaccine for the prevention of recurrent urinary tract infections versus prophylactic treatment with antibiotics. Int. Urogynecol. J. 24, 127–134. 10.1007/s00192-012-1853-5.

78. Lorenzo-Gómez, M.F., Padilla-Fernández, B., García-Cenador, M.B., Virseda-Rodríguez, Á.J., Martín-García, I., Sánchez-Escudero, A., Vicente-Arroyo, M.J., and Mirón-Canelo, J.A. (2015). Comparison of sublingual therapeutic vaccine with antibiotics for the prophylaxis of recurrent urinary tract infections. Front. Cell. Infect. Microbiol. 5, 50. 10.3389/fcimb.2015.00050.

79. Yang, B., and Foley, S. (2018). First experience in the UK of treating women with recurrent urinary tract infections with the bacterial vaccine Uromune(®). BJU Int. 121, 289–292. 10.1111/bju.14067.

80. Nickel, J.C., Saz-Leal, P., and Doiron, R.C. (2020). Could sublingual vaccination be a viable option for the prevention of recurrent urinary tract infection in Canada? A systematic review of the current literature and plans for the future. Can. Urol. Assoc. J. = J. l’Association des Urol. du Canada 14, 281–287. 10.5489/cuaj.6690.

81. Fischer, S., Stegmann, F., Gnanapragassam, V.S., and Lepenies, B. (2022). From structure to function - Ligand recognition by myeloid C-type lectin receptors. Comput. Struct. Biotechnol. J. 20, 5790–5812. 10.1016/j.csbj.2022.10.019.

82. Wells, C.A., Salvage-Jones, J.A., Li, X., Hitchens, K., Butcher, S., Murray, R.Z., Beckhouse, A.G., Lo, Y.-L.-S., Manzanero, S., Cobbold, C., et al. (2008). The macrophage-inducible C-type lectin, mincle, is an essential component of the innate immune response to Candida albicans. J. Immunol. 180, 7404–7413. 10.4049/jimmunol.180.11.7404.

83. Christ, A., Günther, P., Lauterbach, M.A.R., Duewell, P., Biswas, D., Pelka, K., Scholz, C.J., Oosting, M., Haendler, K., Baßler, K., et al. (2018). Western Diet Triggers NLRP3-Dependent Innate Immune Reprogramming. Cell 172, 162–175.e14. 10.1016/j.cell.2017.12.013.

84. Karttunen, J., Sanderson, S., and Shastri, N. (1992). Detection of rare antigen-presenting cells by the lacZ T-cell activation assay suggests an expression cloning strategy for T-cell antigens. Proc. Natl. Acad. Sci. U. S. A. 89, 6020–6024. 10.1073/pnas.89.13.6020.

85. Sancho, D., Mourão-Sá, D., Joffre, O.P., Schulz, O., Rogers, N.C., Pennington, D.J., Carlyle, J.R., and Reis e Sousa, C. (2008). Tumor therapy in mice via antigen targeting to a novel, DC-restricted C-type lectin. J. Clin. Invest. 118, 2098–2110. 10.1172/JCI34584.

86. Sancho, D., Joffre, O.P., Keller, A.M., Rogers, N.C., Martínez, D., Hernanz-Falcón, P., Rosewell, I., and Reis e Sousa, C. (2009). Identification of a dendritic cell receptor that couples sensing of necrosis to immunity. Nature 458, 899–903. 10.1038/nature07750.

87. Garaude, J., Acín-Pérez, R., Martínez-Cano, S., Enamorado, M., Ugolini, M., Nistal-Villán, E., Hervás-Stubbs, S., Pelegrín, P., Sander, L.E., Enríquez, J.A., et al. (2016). Mitochondrial respiratory-chain adaptations in macrophages contribute to antibacterial host defense. Nat. Immunol. 17, 1037–1045. 10.1038/ni.3509.

88. de la Visitación, N., Robles-Vera, I., Toral, M., Gómez-Guzmán, M., Sánchez, M., Moleón, J., González-Correa, C., Martín-Morales, N., O’Valle, F., Jiménez, R., et al. (2021). Gut microbiota contributes to the development of hypertension in a genetic mouse model of systemic lupus erythematosus. Br. J. Pharmacol. 178, 3708–3729. 10.1111/bph.15512.

89. Buenrostro, J.D., Wu, B., Chang, H.Y., and Greenleaf, W.J. (2015). ATAC-seq: A Method for Assaying Chromatin Accessibility Genome-Wide. Curr. Protoc. Mol. Biol. 109, 21.29.1–21.29.9. 10.1002/0471142727.mb2129s109.

90. Kechin, A., Boyarskikh, U., Kel, A., and Filipenko, M. (2017). cutPrimers: A New Tool for Accurate Cutting of Primers from Reads of Targeted Next Generation Sequencing. J. Comput. Biol. a J. Comput. Mol. cell Biol. 24, 1138–1143. 10.1089/cmb.2017.0096.

91. Langmead, B., and Salzberg, S.L. (2012). Fast gapped-read alignment with Bowtie 2. Nat. Methods 9, 357–359. 10.1038/nmeth.1923.

92. Li, H., Handsaker, B., Wysoker, A., Fennell, T., Ruan, J., Homer, N., Marth, G., Abecasis, G., and Durbin, R. (2009). The Sequence Alignment/Map format and SAMtools. Bioinformatics 25, 2078–2079. 10.1093/bioinformatics/btp352.

93. Heinz, S., Benner, C., Spann, N., Bertolino, E., Lin, Y.C., Laslo, P., Cheng, J.X., Murre, C., Singh, H., and Glass, C.K. (2010). Simple combinations of lineage-determining transcription factors prime cis-regulatory elements required for macrophage and B cell identities. Mol. Cell 38, 576–589. 10.1016/j.molcel.2010.05.004.

94. Zhang, Y., Liu, T., Meyer, C.A., Eeckhoute, J., Johnson, D.S., Bernstein, B.E., Nusbaum, C., Myers, R.M., Brown, M., Li, W., et al. (2008). Model-based analysis of ChIP-Seq (MACS). Genome Biol. 9, R137. 10.1186/gb-2008-9-9-r137.

95. Subramanian, A., Tamayo, P., Mootha, V.K., Mukherjee, S., Ebert, B.L., Gillette, M.A., Paulovich, A., Pomeroy, S.L., Golub, T.R., Lander, E.S., et al. (2005). Gene set enrichment analysis: a knowledge-based approach for interpreting genome-wide expression profiles. Proc. Natl. Acad. Sci. U. S. A. 102, 15545–15550. 10.1073/pnas.0506580102.

96. Heger, A., Webber, C., Goodson, M., Ponting, C.P., and Lunter, G. (2013). GAT: a simulation framework for testing the association of genomic intervals. Bioinformatics 29, 2046–2048. 10.1093/bioinformatics/btt343.

97. Ramírez, F., Ryan, D.P., Grüning, B., Bhardwaj, V., Kilpert, F., Richter, A.S., Heyne, S., Dündar, F., and Manke, T. (2016). deepTools2: a next generation web server for deep-sequencing data analysis. Nucleic Acids Res. 44, W160–5. 10.1093/nar/gkw257.

98. Henriksen-Lacey, M., Bramwell, V.W., Christensen, D., Agger, E.-M., Andersen, P., and Perrie, Y. (2010). Liposomes based on dimethyldioctadecylammonium promote a depot effect and enhance immunogenicity of soluble antigen. J. Control. release Off. J. Control. Release Soc. 142, 180–186. 10.1016/j.jconrel.2009.10.022.

